# A novel live cell imaging assay reveals regulation of endosome maturation

**DOI:** 10.1101/2021.06.28.450147

**Authors:** Maria Podinovskaia, Cristina Prescianotto-Baschong, Dominik P. Buser, Anne Spang

## Abstract

Cell-cell communication is an essential process in life, with endosomes acting as key organelles for regulating uptake and secretion of signaling molecules. Endocytosed material is accepted by the sorting endosome where it either is sorted for recycling or remains in the endosome as it matures to be degraded in the lysosome. Investigation of the endosome maturation process has been hampered by the small size and rapid movement of endosomes in most cellular systems. Here, we report an easy versatile live-cell imaging assay to monitor endosome maturation kinetics, which can be applied to a variety of mammalian cell types. Acute ionophore treatment led to enlarged early endosomal compartments that matured into late endosomes and fused with lysosomes to form endolysosomes. Rab5-to-Rab7 conversion and PI(3)P formation and turn over were recapitulated with this assay and could be observed with a standard widefield microscope. We used this approach to show that Snx1- and Rab11-dependent endosomal recycling occurred throughout endosome maturation and was uncoupled from Rab conversion. In contrast, efficient endosomal acidification was dependent on Rab conversion. The assay provides a powerful tool to further unravel various aspects of endosome maturation.

## INTRODUCTION

Endosomes are central organelles in orchestrating cell interactions with the extracellular environment, whether by regulating the composition of signalling molecules at the plasma membrane or by facilitating uptake and digestion of certain nutrients or degrading toxic or no longer needed material. Their wide range of functions is accomplished through a sequential and highly regulated process known as endosome maturation (Huotari and Helenius, 2011; Podinovskaia and Spang, 2018; Spang, 2016). Early endosomes accept incoming cargo from the endocytic vesicles and undergo extensive sorting to package selected cargo into recycling vesicles for the return to the cell surface or to the Golgi, whereas membrane cargo destined for removal is internalised into intraluminal vesicles (ILVs) for its subsequent degradation in endolysosomes. As these sorting endosomes mature into late endosomes, now mainly containing cargo destined for degradation, they no longer accept cargo from the cell surface and acquire properties necessary for their interaction with lysosomes. Upon fusion with lysosomes, late endosomes form endolysosomes, whose highly acidic and hydrolytic milieu facilitates degradation of the remaining cargo and regeneration of the lysosome (Guerra and Bucci, 2016). Throughout this maturation process, the Golgi apparatus supports the endosomal activities by supplying components essential for the progression of endosome maturation, such as proton pump subunits, lysosomal hydrolases, and factors necessary for selective recruitment of GTPases to the endosome (McDermott and Kim, 2015; Nagano et al., 2019).

As endosomes complete their sorting tasks and mature, they undergo extensive changes to their properties to aid their divergent functions. The selective recruitment of GTPases, Rab5 and Rab7 to early and late endosomes, respectively, ensures specificity of interaction with other organelles, such as endocytic vesicles and other early endosomes for Rab5-positive endosomes, and lysosomes for Rab7-positive endosomes (Balderhaar and Ungermann, 2013; Solinger and Spang, 2013). The process of displacement of Rab5 at early endosomes by Rab7 at late endosomes is defined as Rab conversion (Poteryaev et al., 2010; Rink et al., 2005). Additionally, early endosomes contain the signalling lipid PI(3)P, which is further phosphorylated to PI(3,5)P_2_ in late endosomes (Hsu et al., 2015). These lipids serve as organelle identity molecules, facilitating recruitment of components, such as sorting and tethering factors, necessary for endosomal function (Schink et al., 2013). Endosomal acidification is another essential change that must take place for endosomes to mature, with pH ∼6.5, 5.5 and 4.5 characterising early endosomes, late endosomes and lysosomes, respectively (Casey et al., 2010). These changes in GTPase recruitment, PIP composition and acidification status, among others, must be tightly coordinated to ensure unidirectional and aligned adjustments to endosome identity and purpose for the endocytic system to operate properly. However, coordination of such processes during endosome maturation is poorly understood.

A major setback in understanding the kinetics of endosome maturation is lack of experimental systems, which would allow us to monitor endosomes at individual endosome level over prolonged periods of time. The small size of the endosomes and their rapid movement within the cell makes it impractical to track maturing endosomes as they rapidly, within seconds or minutes, move out of field of focus or get lost among other vesicles in the dense perinuclear space. Phagosomes have provided a unique way of studying certain aspects of endosome maturation, allowing for uniform size and synchronisation (Naufer et al., 2018; Podinovskaia et al., 2013). However, these are a specialised subset of endosomes that are involved in minimal amount of sorting and proceed rapidly through endosome maturation, and therefore are not suitable for defining kinetics of classic endosome maturation. Given the present lack of suitable approaches to study endosome maturation, enlarging endosomes might offer a solution to observing individual endosomes over time.

We found that acute nigericin treatment followed by washout led to the formation of enlarged Rab5 positive endosomes that undergo Rab5-to-Rab7 conversion with anticipated kinetics in different cell lines. Other hallmarks of endosome maturation such as PI(3)P, SNX1 and Rab11 recruitment and acidification likewise occurred. Finally, matured late endosomes fused with lysosomes, resulting in functional endolysosomes. Our minimally invasive assay provides an inexpensive and robust way to evaluate relative kinetics of key mediators of endosome maturation at individual endosome level. This assay does not require any specialized equipment, and maturation is detected with the ease of a conventional widefield microscope. We used this assay to investigate whether Rab conversion influences recycling to the plasma membrane and endosomal acidification. We found that recycling to the plasma membrane did not strongly correlate with Rab conversion. Fusing GalT to ratiometric pHlemon (Burgstaller et al., 2019) (GalT-pHlemon) allowed us to follow the degradation pathway to the lysosome and to demonstrate that acidification is slowed down when Rab conversion is blocked, suggesting that Rab conversion is required for efficient acidification during endosome maturation.

## RESULTS

### Short nigericin treatment induces enlarged endosomes that undergo Rab conversion

Endosomes are highly dynamic and motile organelles and, given their small size and frequently indistinct and changing shape, are highly uncooperative to monitoring over extended periods of time by microscopy. The ability to follow dynamic events, such as Rab conversion, at individual endosome level is pivotal for unravelling the mechanisms of endosome maturation. Therefore, we sought a minimally invasive way to enlarge endosomes to make them more distinct and traceable over time. We discovered that a 20-min nigericin treatment of HeLa cells, stably expressing mApple-Rab5 and GFP-Rab7, followed by washout led to the formation of enlarged Rab5- and Rab7-positive endosomes (Fig 1A). Nigericin is an ionophore known to reversibly permeabilise membranes to protons and K^+^ ions. Indeed, the short nigericin treatment disrupted intracellular pH gradient, which re-established within 30 min of washout as visualised by Lysotracker accumulation in treated cells (Fig 1B). The presence of the enlarged Rab5-positive endosomes was often transient but could also last for longer times, whereas enlarged Rab7-positive endosomes persisted until complete recovery of Rab5 and Rab7 morphology by 20 h (Fig 1A). We hypothesised that the enlarged Rab5-positive early endosomes mature to Rab7-positive late endosomes. Therefore, we followed individual endosomes at 1 min intervals, starting from enlarged spherical compartments devoid of either Rab5 or Rab7, and we could indeed observe transient recruitment of Rab5 and its subsequent displacement by Rab7 (Fig 1C, video Fig 1C supplement 1), consistent with previous descriptions of Rab conversion events (Del Conte-Zerial et al., 2008; Poteryaev et al., 2010; Skjeldal et al., 2021). Hence, acute nigericin treatment leads to enlarged compartments that are capable of recruiting Rab5 and undergoing Rab conversion.

**Figure 1.**
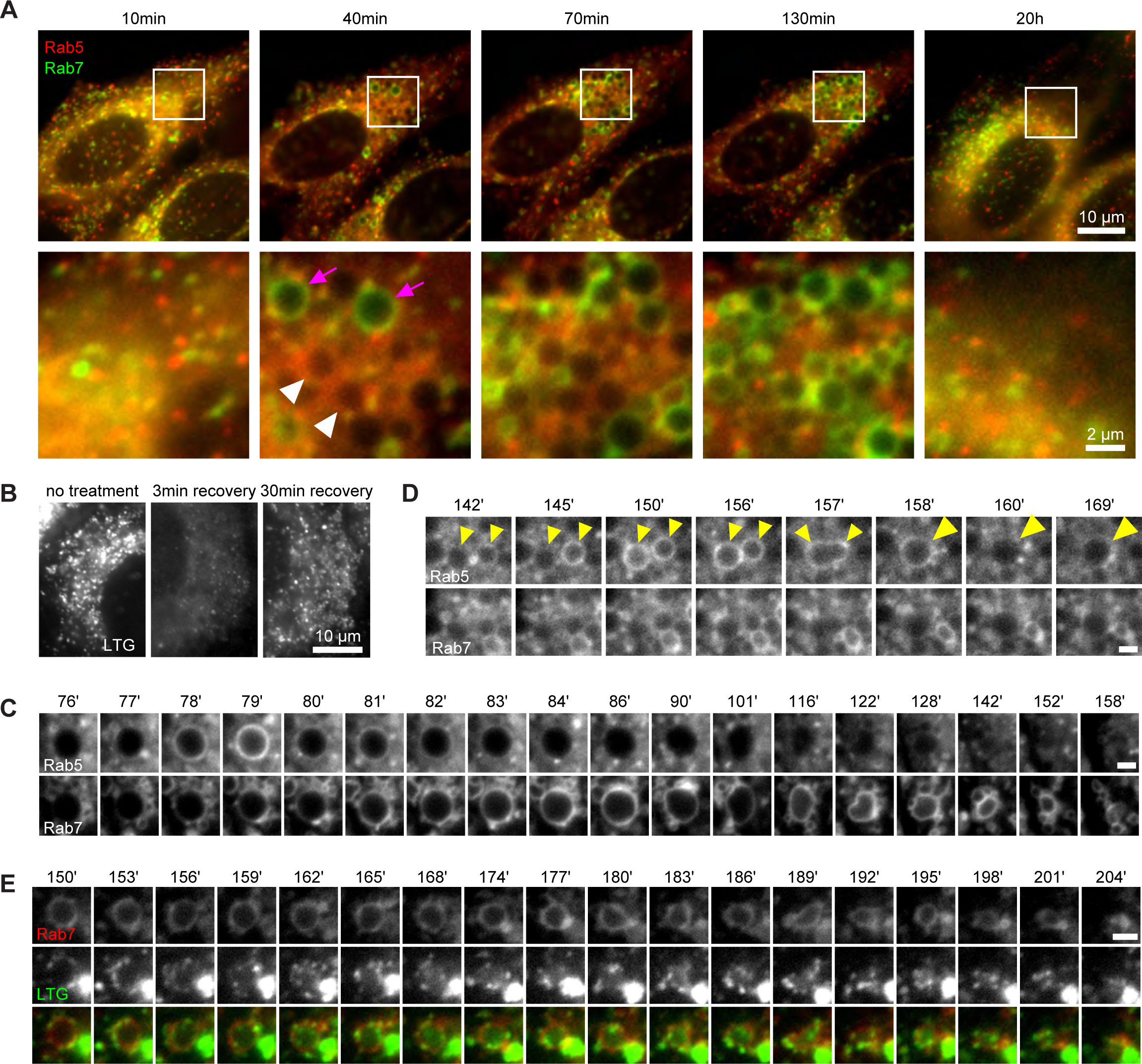
Rab conversion and completion of endolysosomal stages of endosome maturation can be observed in enlarged endosomes, induced with short nigericin treatment. Nigericin was added to HeLa cells at 10 µM for 20 min and washed away, and cells were imaged by time-lapse microscopy. Recovery times are specified relative to removal of nigericin. (A,C,D) Cells stably expressing mApple-Rab5 and GFP-Rab7. (A) Images to show enlarged Rab5- (white arrows) and Rab7- (magenta arrows) positive compartments and return to normal morphology by 20 h. (B) Lysotracker Green (LTG) was added to cells during and following nigericin treatment. Images show rapid re-accumulation of Lysotracker in treated cells. (C) The enlarged endosome was selected to show Rab5 recruitment, Rab conversion and endolysosomal stages of endsome maturation. (D) An example of homotypic fusion of two Rab5-positive endosome and subsequent Rab conversion. (E) Cells stably expressing mApple-Rab7 were imaged in the presence of Lysotracker Green. An enlarged Rab7-positive endosome was selected to show accumulation of Lysotracker concomitant with the loss of spherical shape and a reduction in size of the maturing endolysome. (C-E) Scale bar = 2 µm.

We frequently found that Rab5 recruitment was initiated after a Rab5-positive endocytic vesicle or endosome was touching or fusing with an enlarged compartment (Fig 1C; Fig 1, figure supplement 1A), suggesting either kiss-and-run or fusion with a Rab5-positive structure drove Rab5 recruitment. Therefore, endocytic events from the plasma membrane are also required to form enlarged Rab5 positive early endosomes. We also observed homotypic Rab5 endosomal fusions, a hallmark of early endosomes, indicating that the enlarged Rab5-positive structures behaved as bona fide early endosomes (Fig 1D, video Fig 1D supplement 1). Following Rab conversion, the spherical Rab7-positive endosomes persisted over a range of several minutes to several hours and remained Lysotracker-negative, with lysosomes seen as Lysotracker-positive puncta circling around the endosomes. Once the endosome acidified sufficiently to accumulate Lysotracker, it lost its spherical shape and got torn apart until no longer detectable (Fig 1C and E, video Fig 1E supplement 1), consistent with endolysosomal stages of endosome maturation. Fusion of the enlarged endosomes with Dextran-AF488-loaded lysosomes was apparent through accumulation of Dextran-AF488 in the enlarged endosomes (Fig 1, figure supplement 1B). Thus, acute nigericin treatment could induce the formation of large early endosomes, which could be observed to mature into late endosomes, and subsequently fuse with lysosomes and undergo endolysosome-to-lysosome maturation. This acute treatment may provide the basis of a powerful assay that could be employed to follow individual maturing endosomes.

How common is this phenomenon of the enlarged endosome induction by acute pharmaceutical treatment? First, we checked whether acute treatment with another ionophore, monensin, or the weak base NH_4_Cl, which perturbs the pH gradient, would have a similar effect. Indeed, we observed transient Rab5 recruitment and the more extended Rab7 recruitment at the enlarged endosomes of NH_4_Cl pre-treated cells, and likewise, gradual Rab7 recruitment following acute monensin treatment (Fig 1, figure supplement 2). Therefore, interfering with ion homeostasis and membrane potential appear to solicit a similar stress response as nigericin, resulting in formation of enlarged endosomes. Second, we investigated whether this effect was cell line specific or more generally applicable. We tested the epithelial cell line HEK293, the fibroblast-like cell line COS1 and the neuronal line Neuro2. In all three cell lines we could observe enlarged endosomes that were either Rab5 or Rab7 positive (Fig 1, figure supplement 3). Therefore, enlarged endosome induction is not restricted to nigericin treatment of HeLa cells but rather is applicable to a wide range of experimental systems.

### TGN membranes transition into endosomes after acute nigericin treatment

Short nigericin treatment led to enlarged endosomes, however, they did not start out as Rab5-positive entities (Fig 1C). Therefore, we investigated the origin of the membranes for these compartments. Electron microscopy images revealed that the enlarged compartments originate at the trans face of the Golgi (Fig 2A), in line with previous reports of ionophore treatment leading to the swelling of the trans-Golgi leaflet (Ledger et al., 1980; Morre et al., 1983). We ruled out contribution from the autophagy pathway by staining mApple-Rab5 expressing cells with LC3b antibody and showing no detectable autophagy induction or LC3b presence at the enlarged endosomes at 60 min post nigericin treatment (Fig.2, figure supplement 1A). To determine whether the swollen TGN membranes would enter the endosomal pathway, we performed immuno-electron microscopy with HeLa cells stably expressing trans-Golgi marker GalT-GFP after acute nigericin treatment. The micrographs demonstrate the presence of GalT in the enlarged trans-Golgi network (TGN) compartments and in ILVs of multivesicular bodies at later time points (Fig 2B). Therefore, the membranes that acquire Rab5 and convert to Rab7 positive endosomes are probably derived from the TGN. This swelling of the TGN is likely a transient response to the acute stress because after 48 h the Golgi had recovered from the treatment (Fig 2C). Consistent with this notion, we occasionally observed swollen Golgi leaflets also in untreated cells signifying a process that occurs naturally in the cell, which we are uniquely amplifying with acute perturbation (Fig 2A). Indeed, the cells continued to grow and divide (Fig.2, figure supplement 1B), and after an initial slow start, the nigericin-treated cells recover their doubling rate within 24 h (Fig.2, figure supplement 1C). In line with previous reports (Merion and Sly, 1983), our findings indicate that short nigericin treatment induces reversible changes and has minimal impact on cell health.

**Figure 2.**
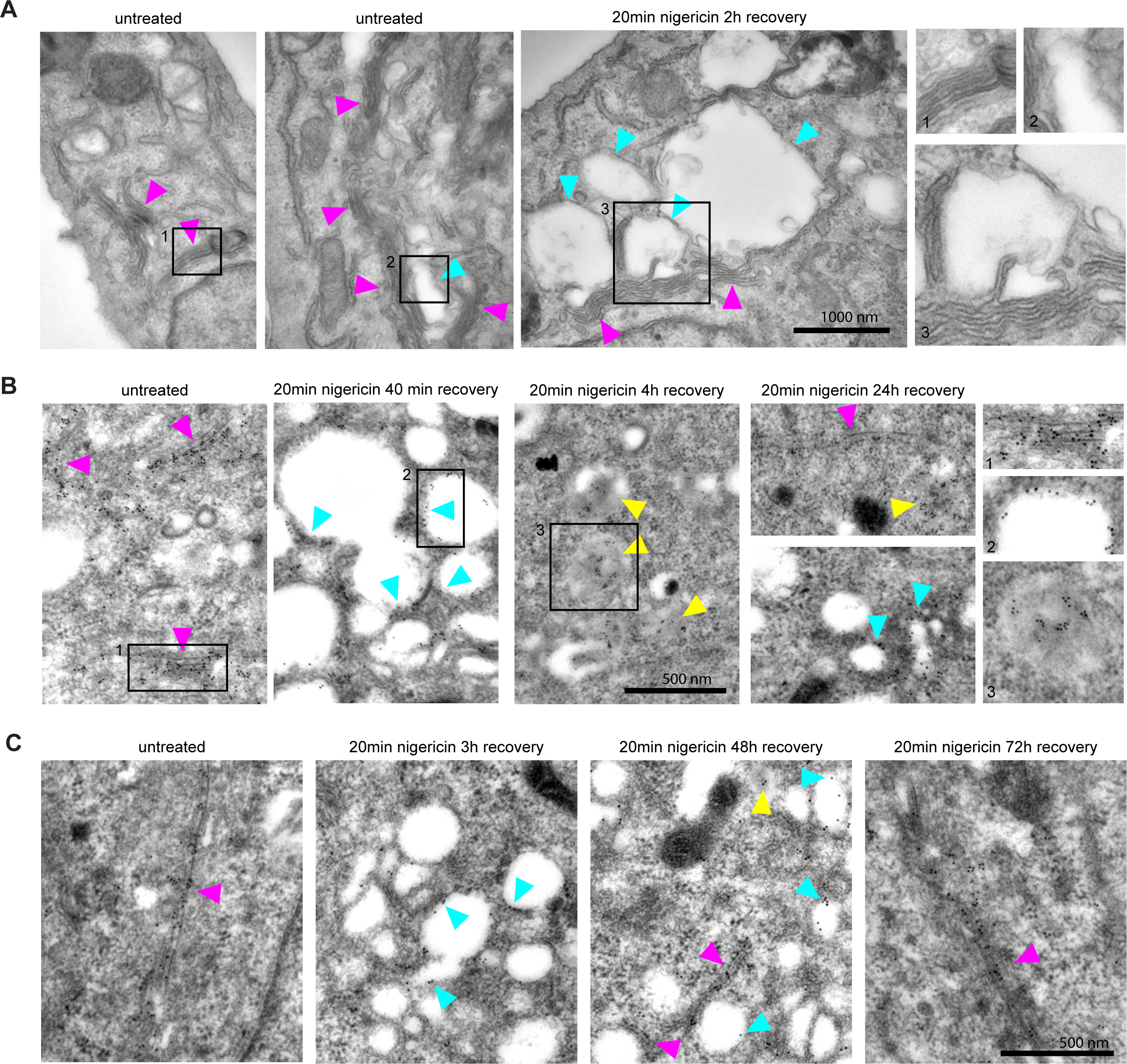
Nigericin-induced enlarged compartments originate at the Golgi and contain trans-Golgi marker GalT, later found in ILVs, with most enlarged compartments resolved by 48 h. Nigericin was added to HeLa cells for 20 min and washed away, and cells were processed for electron microscopy (A-C), imaged by time-lapse microscopy (D) or harvested for counting (E) at specified times after the wash. (A) Cells stained with osmium tetroxide and potassium hexacyanoferrate reveal large spherical compartments (cyan arrows) originating at the trans-face of the Golgi (magenta arrows) in nigericin-treated cells and, occasionally, in untreated cells. (B,C) Cells stably expressing GalT-GFP were stained with anti-GFP and 12 nM immuno-Gold to reveal GalT-GFP at the Golgi (magenta arrows), the limiting membrane of the enlarged compartments (cyan arrows) as well as in ILVs of the enlarged MVBs at later time points (yellow arrows).

To corroborate our results, we monitored HeLa cells stably expressing GalT-GFP by fluorescence microscopy following acute nigericin treatment. Golgi vesiculation was observed within 15 min of nigericin washout (Fig 3A, video Fig 3A supplement 1). We observed similar results when we used monensin as ionophore (Fig.2, figure supplement 1D). A large fraction of these vesicles would adopt early endosomal identity because individual GalT-positive structures acquired Rab5 over time (Fig 3A), as also observed by immuno-electron microscopy (Fig 3B). Moreover, similar to the transiently transfected GalT-GFP, endogenous GalT persisted in the endosomes (Fig 3C), consistent with the observations of its subsequent internalisation into ILVs (Fig 2B and 10D), and Golgi morphology was fully recovered within 48h (Fig.2, figure supplement 1E). The contribution of cargo from the endocytic pathway to the enlarged compartments was evidenced by the addition of Dextran-AF488 for 1 h to the cell medium of nigericin-treated cells and its detection in the Rab5-positive enlarged endosomes after washing the dextran away (Fig.2, figure supplement 1F). Taken together, our findings suggest that acute nigericin treatment leads to enlarged Golgi-derived compartments that are able to acquire early endosomal identity and mature into late endosomes. Thereby, acute nigericin treatment provides us with a means to generate functionally competent enlarged endosomes that can be monitored at individual endosome level by widefield microscopy over extended periods of time to define the kinetics of a wide range of mediators of endosome maturation.

**Figure 3.**
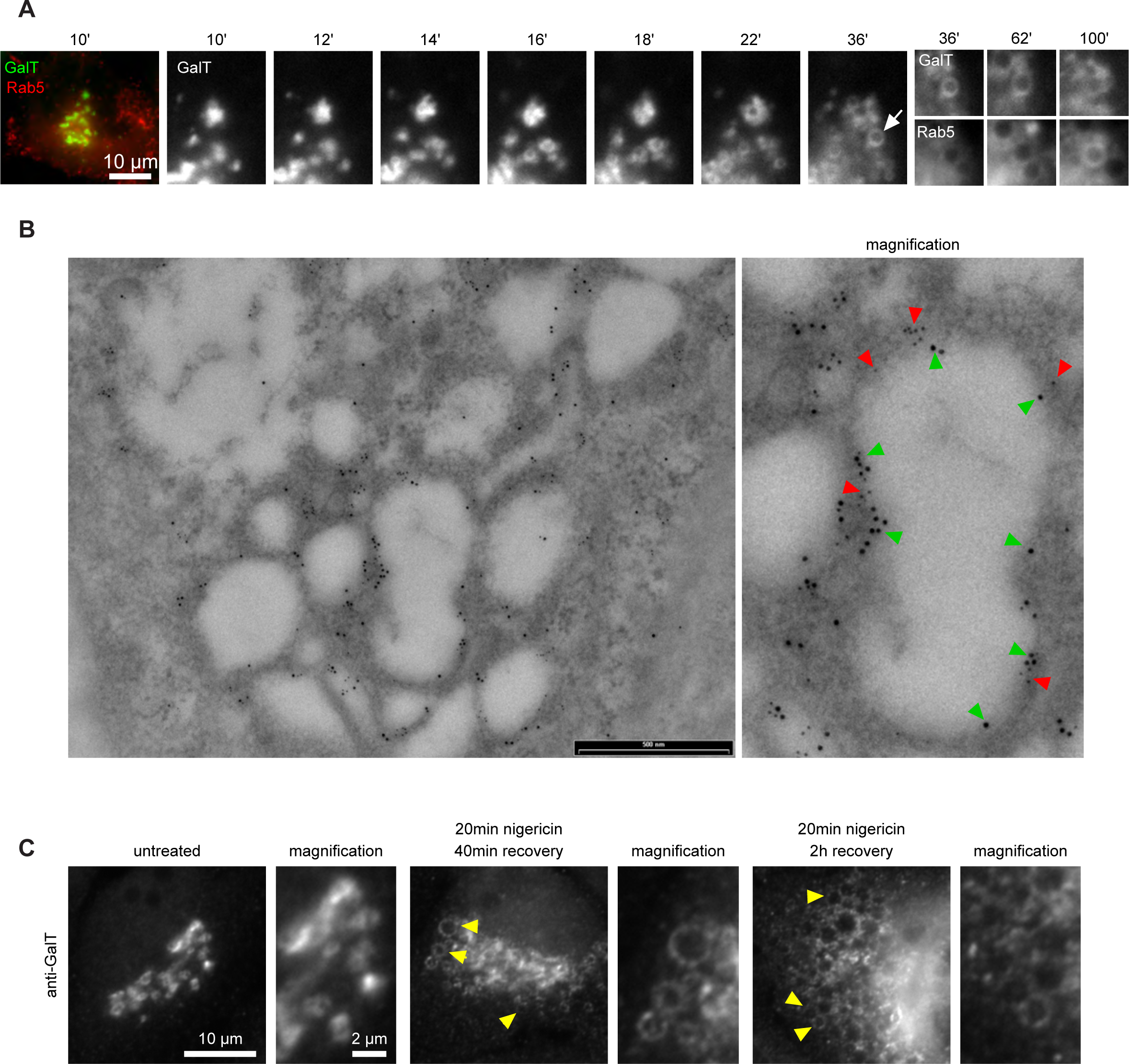
Short nigericin treatment leads to trans-Golgi vesiculation and subsequent acquisition of Rab5. Nigericin was added to HeLa cells for 20 min and washed away, and cells were imaged by time-lapse microscopy (A), processed for electron microscopy (B) or for immunofluorescence (C) at specified times after the wash. (A,B) HeLa cells stably expressing GalT-GFP and transiently transfected with mApple-Rab5. (A) Representative kinetic of Golgi vesiculation post nigericin treatment as visualised with GalT-GFP. The selected vesiculated compartment (arrow), initially negative for Rab5 subsequently becomes positive for both markers. (B) Immuno-EM image of cells at 2 h post recovery, with 12 nm Gold labelled GFP (green arrows) and 5 nm Gold labelled mApple (red arrows) present at the enlarged compartments. Scale bar = 500 nm. (C) Images of cells stained with anti-GalT to reveal endogenous GalT presence at the enlarged compartments.

### Rab conversion occurs with anticipated kinetics on enlarged endosomes

Having established a novel assay to study endosome maturation, we used it first to revisit the kinetics of Rab conversion. The formation of enlarged early endosomes was asynchronous and therefore we imaged over several hours without significant loss of fluorescence signal. We captured many events of Rab conversion for further analysis and signal quantification. Endosomes that were initially negative for Rab5 and acquired Rab5 during the time course were chosen for analysis. For quantification purposes, we measured the mean fluorescence intensity of the rim of the enlarged endosome at all time points when the endosome was detectable (Fig 4A). Following transient Rab5 recruitment, all selected endosomes underwent Rab conversion. Initially Rab5 was recruited uniformly to the rim of the endosome, but could segregate also into distinct domains, before becoming completely displaced by Rab7 (Fig 4B, Fig. 4, figure supplement 1A). Consistent with previous findings, Rab5 levels dropped when Rab7 reached about 50% of its maximal level (Fig. 4B-D) (Del Conte-Zerial et al., 2008; Poteryaev et al., 2010; van der Schaar et al., 2008). Moreover, Rab conversion was completed within 4 min after its initiation, which is similar to previously reported observations (Del Conte-Zerial et al., 2008; Poteryaev et al., 2010; Rink et al., 2005). Once Rab5 was fully removed, Rab7 plateaued off showing stable presence at the late endosome (Fig 4D; Fig. 4, figure supplement 1B). Occasionally, Rab5 produced multiple peaks, with Rab7 plateauing off after the latest Rab5 peak (Fig. 4, figure supplement 1C and D). Such Rab5 behaviour may indicate the reversible nature of endosome maturation and existence of checkpoints to ensure alignment of parallel processes. Our results closely agree with Rab conversion kinetics in other systems and further refine Rab conversion kinetics in human cells. Empowered by this strict sequential kinetics of Rab5 and Rab7 in maturing endosomes, we next explored the kinetics of other mediators of endosome maturation relative to either Rab5 or Rab7 recruitment, using the maximum peak of Rab5 or the 50% of the maximal fluorescence intensity of Rab7 as reference point for Rab conversion.

**Figure 4.**
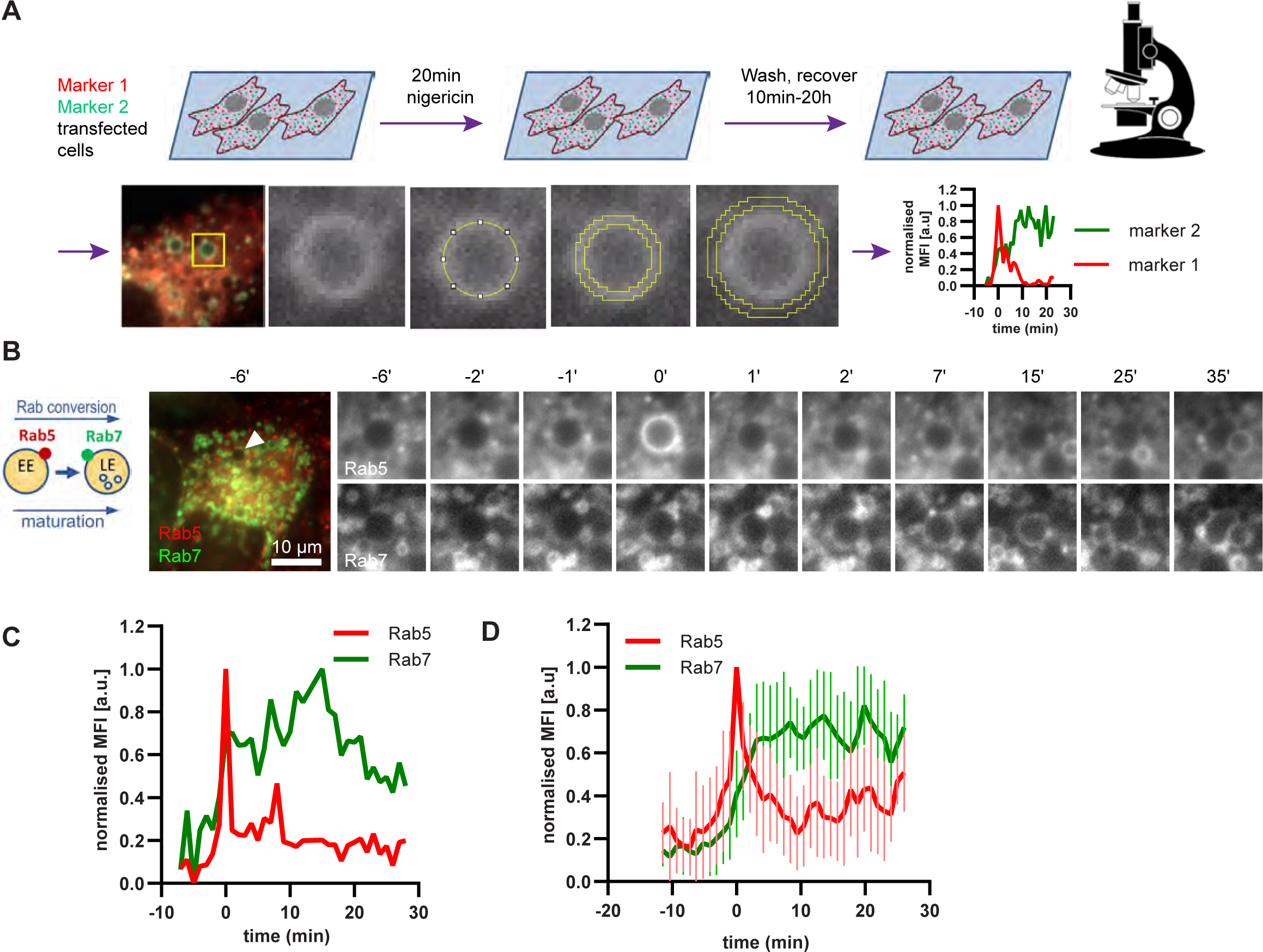
Enlarged endosomes recruit Rab5 and undergo Rab conversion with anticipated kinetics. (A) Scheme to show experimental flow, starting with transfection of cells of choice with selected markers, followed by short nigericin treatment, and time-lapse microscopy during the recovery phase, with subsequent quantification of mean fluorescence intensity (MFI) of the chosen markers at the rim of the enlarged endosomes, and the resulting kinetic plots of background-subtracted MFI normalised for maximum and minimum values over the entire time course of the endosome. Since endosome maturation is asynchronous, relative time is calculated by using Rab5 peak as a reference for Rab conversion and set to t=0. The plot shown in the scheme represents the kinetic of the images in Figure 1C (marker 1 as Rab5 and marker 2 as Rab7). (B,C,D) HeLa cells, stably expressing mApple-Rab5 and GFP-Rab7 were treated for 20 min with nigericin, washed and imaged over a 3 h period. (B) Time-lapse images of a representative endosome to show transient Rab5 recruitment and its subsequent displacement by Rab7. (C) Corresponding graph of MFI of Rab5 and Rab7 at the rim of the endosome in (B) during and around the time of Rab conversion. Numerical data for all analyzed endosomes is available in Figure 4D - Source Data 1. (D) Averaged Rab5 and Rab7 kinetics of 27 endosomes. Error bars represent standard deviation. Representative graph of three independent experiments.

### PI(3)P levels peak concomitantly with Rab5 levels

Driving early endosome identity, Rab5 recruits the PI(3)P kinase VPS34 and forms a positive feedback loop with PI(3)P (Zerial and McBride, 2001). Coincidence detection of PI(3)P levels and the Rab5GEF Rabex5 by the Rab7 GEF Mon1/CCZ1 was proposed to drive Rab conversion and endosome maturation (Poteryaev et al., 2010) and subsequently trigger the formation of PI(3,5)P2 (Compton et al., 2016; Dove et al., 2009). We analyzed cells expressing mApple-Rab5 and the PI(3)P marker GFP-FYVE. As expected, Rab5 and PI(3)P appeared concomitantly on enlarged early endosomes (Fig 5, video Fig 5A supplement 1). However, after Rab5 peaked, we observed a slight delay in the disappearance of GFP-FYVE (Fig. 5C, Fig. 5, figure supplement 1), suggesting that the onset of PI(3)P to PI(3,5)P2 conversion occurs with some delay. Nevertheless, our data are consistent with a tight temporal and spatial regulation of PI(3)P levels on endosomes during maturation.

**Figure 5.**
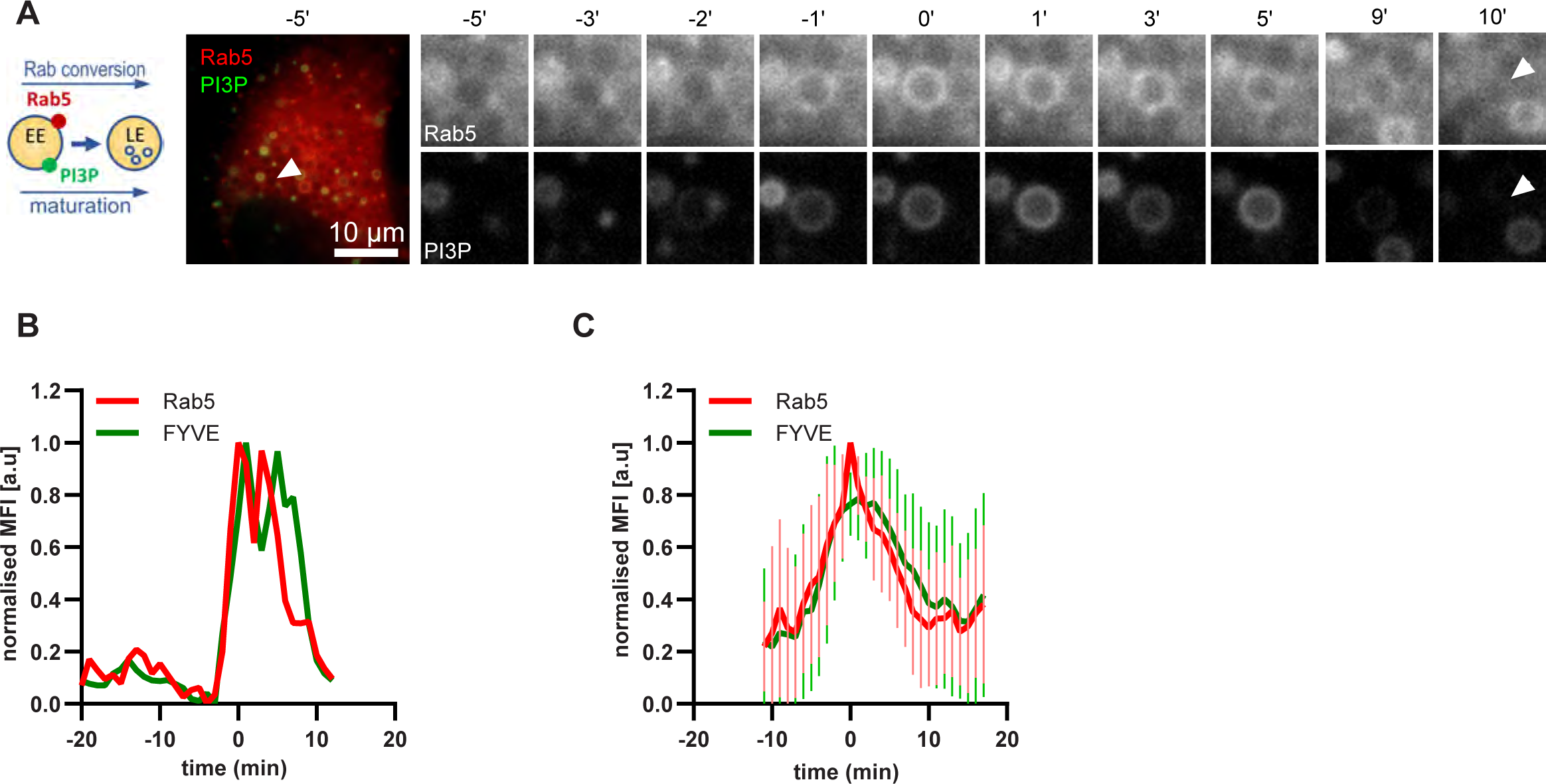
PI(3)P is recruited to endosomes concomitantly with Rab5. HeLa cells, stably expressing mApple-Rab5 and transiently transfected with the PI(3)P marker, GFP-FYVE, were treated for 20 min with nigericin, washed and imaged over 3 h, as described in Figure 4A. (A) Time-lapse images of a representative endosome to show transient Rab5 recruitment accompanied by PI(3)P. (B) Corresponding graph of normalised mean fluorescence intensity of Rab5 and FYVE at the rim of the endosome in (A) over the time the endosome was detectable. (C) Averaged Rab5 and PI(3)P kinetics of 19 endosomes. Error bars represent standard deviation. Representative graph of three independent experiments. Numerical data for all analyzed endosomes is available in Figure 5C - Source Data 1.

### Snx1 recruitment is initiated with Rab5 recruitment and can persist during Rab7 stages

A major property of endosomes is their ability to undergo extensive sorting, to recycle components back to the cell surface and to the Golgi and to internalise membrane cargo destined for ILV-mediated degradation. Our electron microscopy data provides evidence that the nigericin-induced enlarged endosomes are capable of ILV formation and internalisation of the GalT marker (Fig 2B). Additionally, the presence of Snx1-GFP in transient punctate microdomains or tubular protrusions at the enlarged endosomes suggests active sorting from the endosome to the plasma membrane and the Golgi (Fig 6A). To determine whether the Snx1-mediated sorting is coordinated with Rab conversion, we analyzed cells co-expressing Snx1-GFP and mApple-Rab5 (Fig 6A), and recorded the presence of Snx1 at the enlarged endosomes as they acquired and removed Rab5. Snx1 assembly on endosomes occurred concomitantly with Rab5 recruitment (Fig 6 A-C; video Fig 6A supplement 1; Fig. 6, figure supplement 1 A and B), pointing to a potential coordination. Snx1 assembly at the endosomes was highly dynamic, forming one or multiple domains at a time (Fig. 6, figure supplement 1 C). We observed weak correlation of Snx1 and Rab5 localization in discrete domains on early endosomes (Fig 6D and E). Moreover, Snx1 levels either declined during Rab conversion or persisted for a while. Our data indicate that Snx1 recruitment on early endosomes occurs simultaneously with Rab5, but that Snx1 microdomains could either co-exist with or exist independently of Rab5, suggesting that Rab5 may promote Snx1 recruitment but is not essential for its maintenance or dynamics at endosomes.

**Figure 6.**
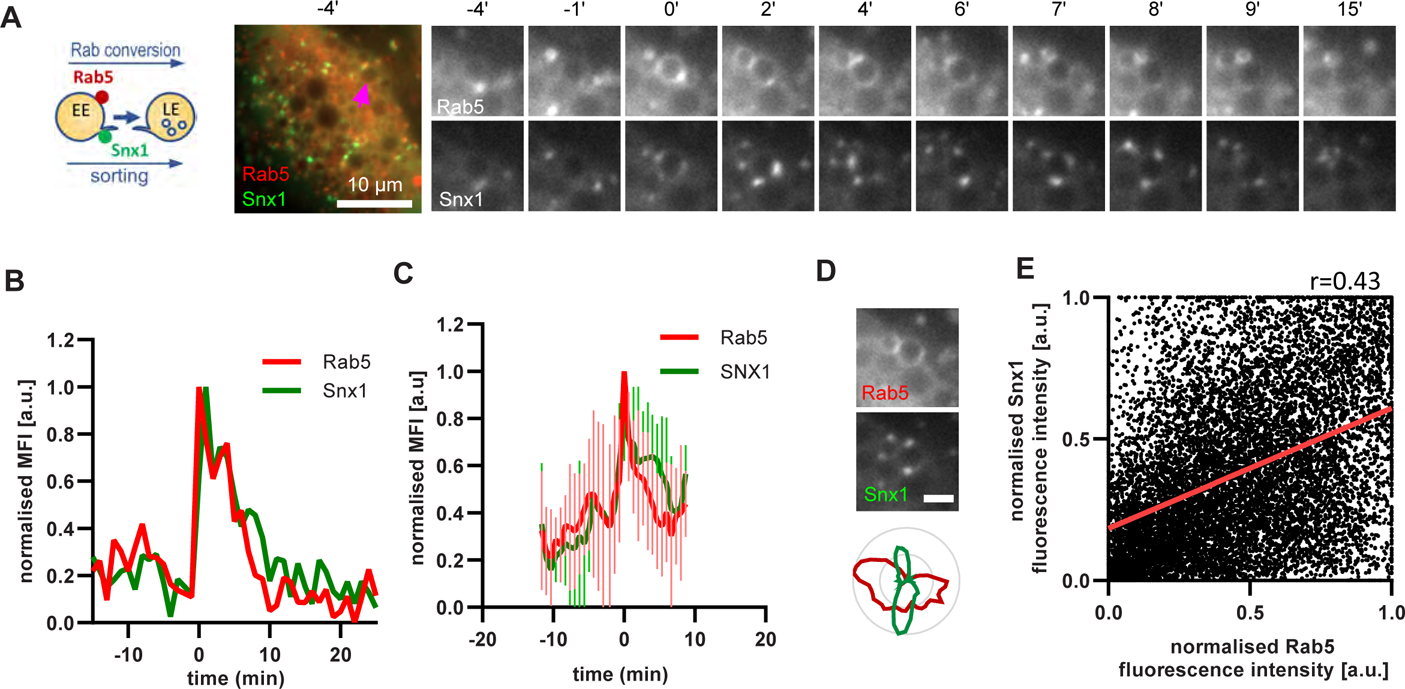
Snx1 subdomain formation at the endosomes initiates with Rab5 recruitment and peaks during Rab conversion stages. HeLa cells, stably expressing mApple-Rab5 and transiently transfected with Snx1-GFP, were treated for 20 min with nigericin, washed and imaged over 3 h, as described in Figure 4A. (A) Time-lapse images of a representative endosome to show Snx1 subdomain formation relative to Rab5 recruitment. (B) Corresponding graph of normalised mean fluorescence intensity (MFI) of Rab5 and Snx1 at the rim of the endosome in (A) over the time the endosome was detectable. (C) Averaged Rab5 and Snx1 kinetics of 12 endosomes. Error bars represent standard deviation. Representative graph of three independent experiments. Numerical data for all analyzed endosomes is available in Figure 6C - Source Data 1. (D) Images of Rab5 and Snx1 at an enlarged endosome and a corresponding line profile of normalised fluorescence intensity along the rim to show co-existence as well as independence of subdomains of the two markers. Scale bar = 2 µm. (E) Correlation plot of normalised fluorescence intensity of Rab5 and Snx1 as measured in (D) for 14 endosomes for a total of 118 time points, and a corresponding regression line. Pearson’s correlation r=0.43. Pooled data from two independent experiments. Numerical data for all analyzed endosomes is available in Figure 6E - Source Data 1.

To corroborate the apparent lack of strict coordination between Rab5 removal and Snx1 persistence at the endosomes, we co-expressed Snx1-GFP and mApple-Rab7 (Fig 7A, video Fig 7A supplement 1). Consistent with Snx1 presence during the Rab5 phase, Snx1 recruitment peaked during early stages of Rab7 recruitment, when the endosome is expected to be Rab5-positive (Fig 7B-D). In about one third of all Rab7-positive endosomes analysed, Snx1 recruitment was transient and was no longer present after Rab7 peaked or levelled off to indicate completed Rab conversion (Fig 7B and E). In another third of analysed endosomes, Snx1 initially displayed the same kinetics, but was recruited back again to the late Rab7-positive endosome (Fig 7C and F), suggesting that Rab5 may be dispensable for Snx1 recruitment to late endosomes. In the remaining subset of endosomes, Snx1 peaked and persisted throughout endosome maturation (Fig 7D and G). Analysis of Rab7 and Snx1 domains at the endosome again revealed weak correlation of Rab7 and Snx1 domains (Fig. 6, figure supplement 1D and E). Taken together, our data suggest that sorting into recycling pathways is most likely initiated very early on endosomes but can also persist on late endosomes, indicating a continuous process independent of Rab conversion.

**Figure 7.**
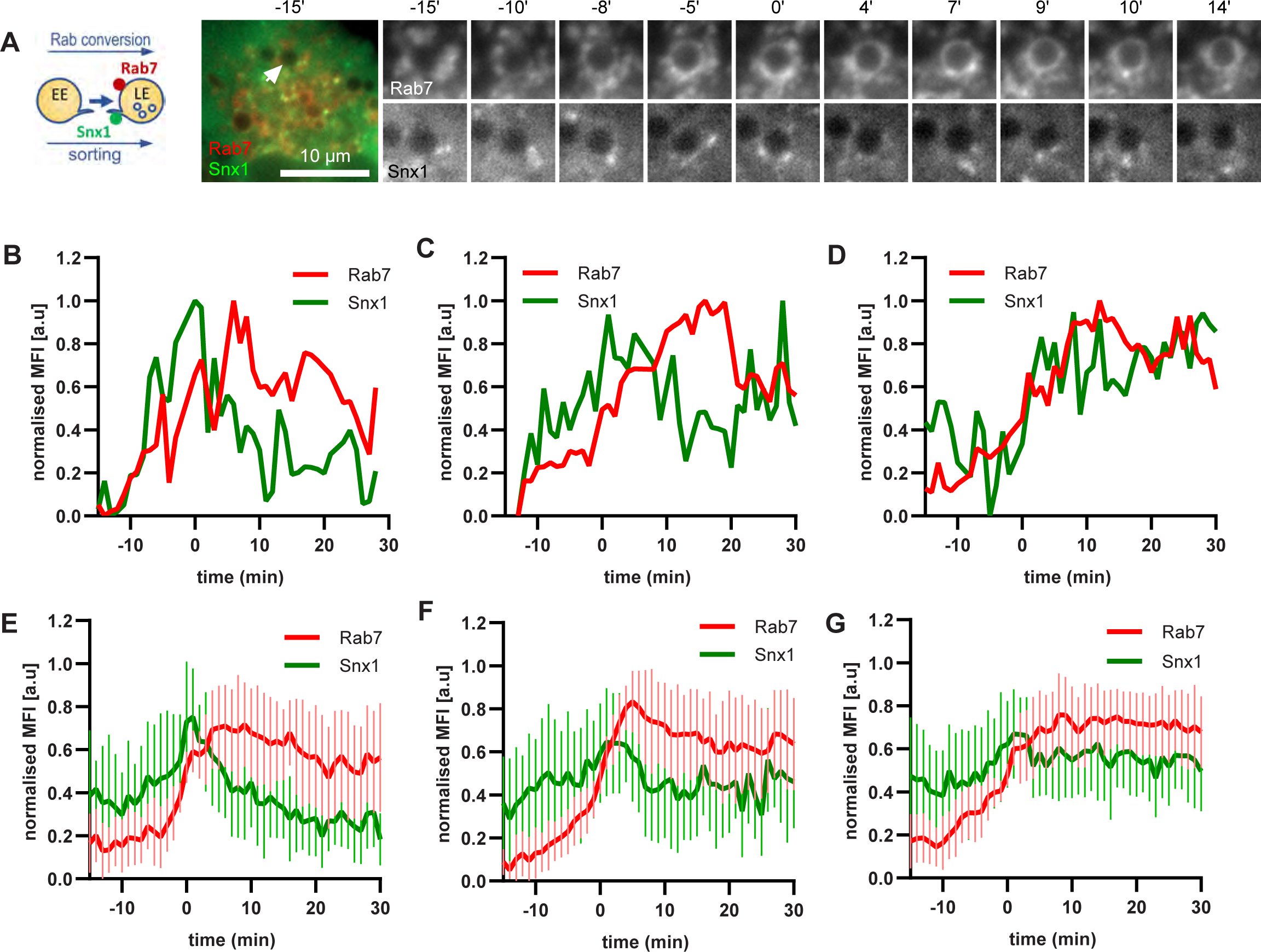
Snx1 subdomain formation at the endosomes initiates with Rab5 recruitment, peaks during Rab conversion stages and continues in late endosomes. HeLa cells, stably expressing mApple-Rab7 and transiently transfected with Snx1-GFP, were treated for 20 min with nigericin, washed and imaged over 3 h, as described in Figure 4A. (A) Time-lapse images of a representative endosome to show Snx1 subdomain formation relative to and Rab7 recruitment. (B) Corresponding graph of MFI of Rab7 and Snx1 at the rim of the endosome in (A) over the time the endosome was detectable, to show Snx1 peaking during Rab conversion. (C,D) Additional graphs of MFI of Rab7 and Snx1 at the rim of endosomes to show the second Snx1 peak (C) or continuing Snx1 presence (D). (E,F,G)) Averaged Rab7 and Snx1 kinetics binned into the three patterns of Snx1 recruitment as observed in (B,C,D), representing 19, 21 and 20 endosomes for the single peak, double peak and continuing presence of Snx1, respectively. Error bars represent standard deviation. Three independent experiments were performed, and data pooled. Numerical data for all analyzed endosomes is available in Figure 7 - Source Data 1.

### Interaction of early and late endosomes with Rab11 proceeds independently of Rab5 or Rab7

To further support our hypothesis that sorting can occur continuously from early to late endosomes, we examined the behaviour of a key component of the recycling pathway to the plasma membrane, Rab11. If our hypothesis were correct, we would expect Rab11 to show similar behaviour as Snx1. Therefore, we investigated the dynamics of GFP-Rab11 in relation to Rab5 or Rab7 positive endosomes. Rab11 docks on the tubular part of maturing/sorting endosomes and promotes the recycling of cargo to the plasma membrane (Solinger et al., 2020; van Weering et al., 2012). Surprisingly, Rab11 positive vesicles contacted early endosomes even before strong Rab5 recruitment (Fig 8A-C, video Fig 8A supplement 1). Moreover, Rab11 interaction with endosomes appeared to be independent of Rab5, Rab7, or Rab conversion as it continued throughout endosome maturation (Fig. 8, video Fig 8E supplement 1; Fig. 8, figure supplement 1). Thus, our data indicate that, similar to sorting, recycling to the plasma membrane appears to be largely decoupled from Rab conversion.

**Figure 8.**
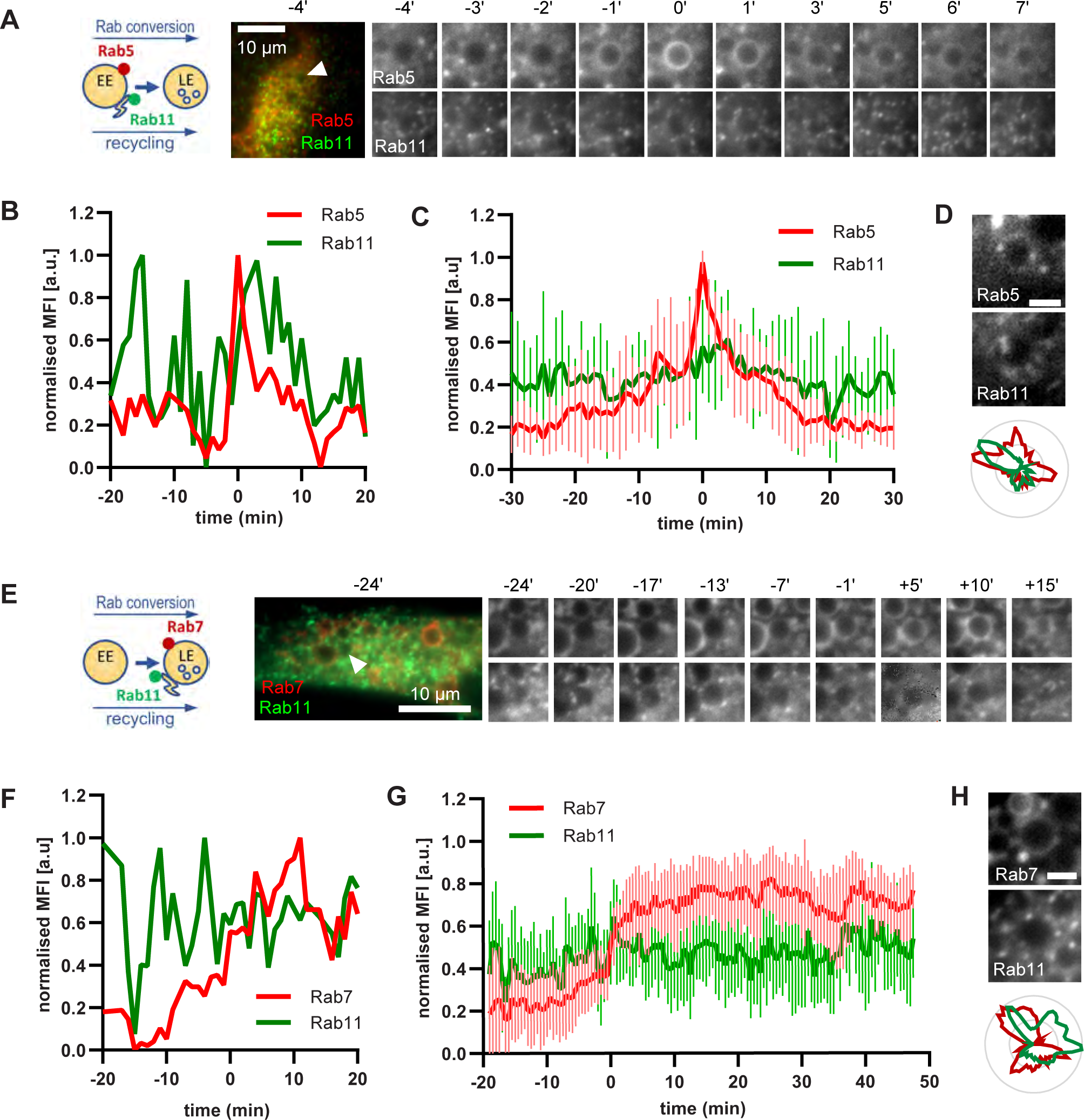
Rab11 interacts with the maturing endosome independently of Rab5 or Rab7. HeLa cells, stably expressing mApple-Rab5 (A-D) or mApple-Rab7 (E-H) and transiently transfected with GFP-Rab11, were treated for 20 min with nigericin, washed and imaged over 3 h, as described in Figure 4A. (A,E) Time-lapse images of a representative endosome to show continuous Rab11 interaction with the maturing endosome relative to Rab5 (A) or Rab7 (E) recruitment. (B,F) Corresponding graphs of normalised mean fluorescence intensity of Rab5 (B) or Rab7 (F) and Rab11 at the rim of the endosome in (A) or (E), respectively, over the time the endosome was detectable. (C,G) Averaged Rab5 (C) or Rab7 (G) and Snx1 kinetics of 16 and 15 endosomes, respectively. Error bars represent standard deviation. Representative graphs each of three independent experiments. Numerical data for all analyzed endosomes is available in Figure 8C - Source Data 1 and Figure 8G - Source Data 1. (D,H) Images of Rab5 (D) or Rab7 (H) and Rab11 at an enlarged endosome and corresponding line profiles of normalised fluorescence intensity along the rim to show co-existence as well as independence of subdomains of Rab11 and the two markers. Scale bar = 2 µm. Numerical data for analyzed endosomes is available in Figure 8D and S8 - Source Data 1 and Figure 8H and S8 - Source data 1.

### GalT-pHlemon is a reliable reporter for pH measurements along the endocytic pathway

Both, Rab conversion and acidification are essential for endosome maturation but how the two are coordinated is poorly understood. To follow endosomal acidification at individual endosome level, we have tested several available endosomal pH sensors, such as mApple-Lamp1-pHluorin and NHE6-pHluorin2 (Ma et al., 2017), however, we found that these sensors were predominantly retained in the ER following transient transfection in our system. We exchanged the pHluorin tag on Lamp1 and NHE6 with the recently developed pH-responsive ratiometric probe, pHlemon (Burgstaller et al., 2019), but this did not improve the export from the ER. Therefore, we replaced the GFP of the GalT-GFP construct with pHlemon. Since GalT was present at the Golgi-derived enlarged compartments, and later in Rab5-positive endosomes and ILVs (Fig 2B, 3A-B), we hypothesised that the sensor anchored to GalT will illuminate the entire endosome maturation pathway, from early endosomes to endolysosomes. The pHlemon probe consists of yellow and mTurquoise2 fluorescent proteins in tandem, with YFP reducing and mTurquoise2 slightly increasing fluorescence upon acidification in a reversible manner (Burgstaller et al., 2019). Untreated cells expressing the GalT-pHlemon sensor displayed a characteristic Golgi-ribbon appearance in both YFP and CFP channels as well as punctate appearance of CFP signal alone, indicative of highly acidified lysosomes or endolysosomes (Fig 9A). We could reliably detect YFP/CFP ratios over the pH 4.0-7.5 range (Fig 9B, Fig. 9, figure supplement 1), allowing for accurate pH measurements of the entire endolysosomal pathway. Our sensor designated pH 6.2 for the Golgi-ribbon structures and pH 4.0-5.7 for the lysosomes and endolysosomes, as detected by the CFP puncta, in untreated cells (Fig 9C). Most importantly, our sensor located to the nigericin-induced enlarged endosomes and indicated a pH range between 5.5 and 6.6 at 50 min washout, reflective of the different stages of maturation (Fig 9D, video Fig 9D supplement 1). Therefore, GalT-pHlemon is a useful tool to read-out pH in the endosomal system.

**Figure 9.**
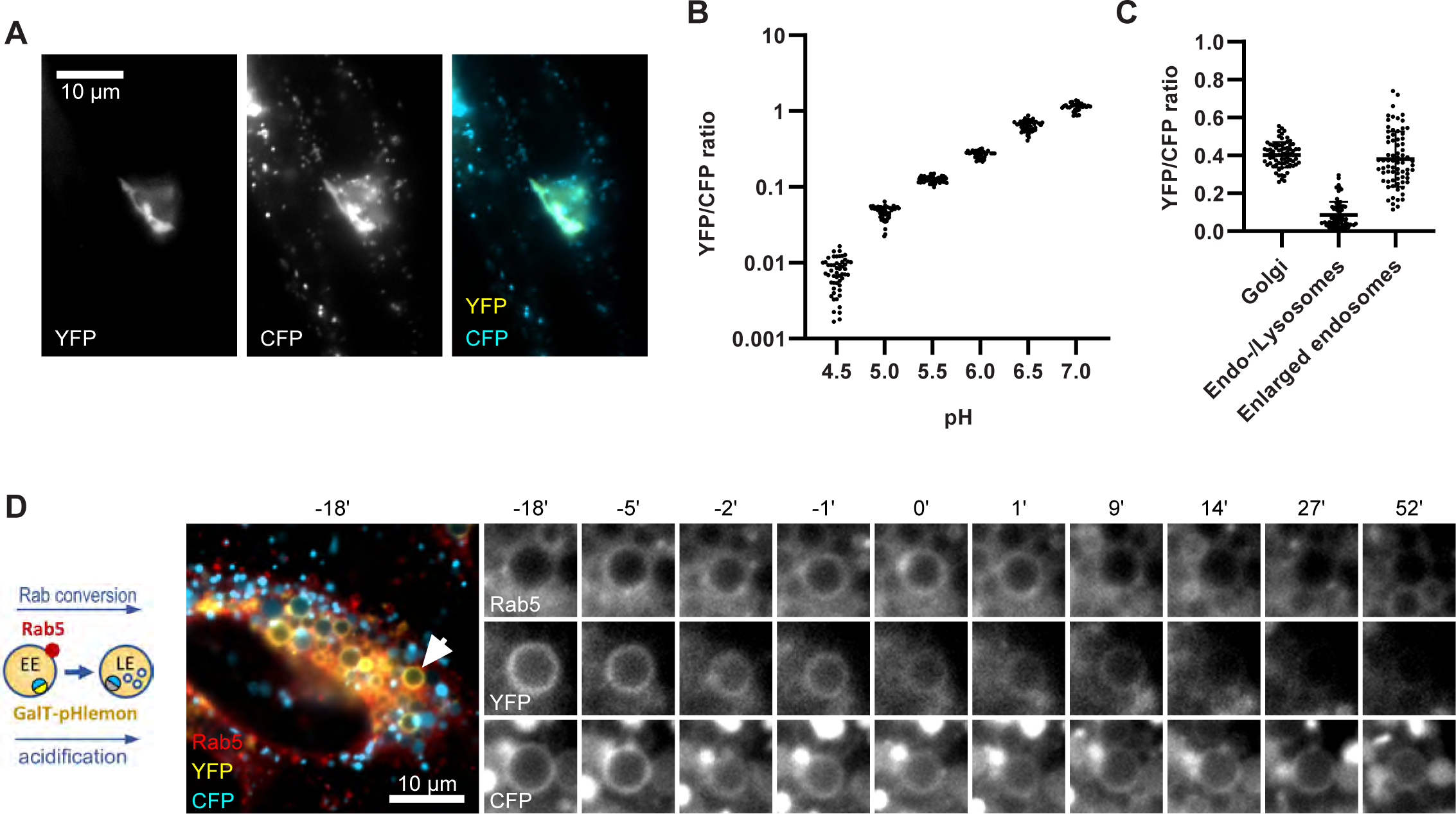
GalT-pHlemon sensor detects endosomal acidification. (A-C) HeLa cells were transiently transfected with the ratiometric pH sensor, GalT-pHlemon. (A) Images of cells to show Golgi-ribbon distribution of GalT-pHlemon in both YFP and CFP channels as well as cytosolic CFP-filled puncta in CFP channel only, representing highly acidified organelles. (B) Graph to show robust response of GalT-pHlemon sensor to pH 4.5-7.0 range as displayed by YFP/CFP ratio measurements in cells incubated with calibration buffers of specified pH values. Numerical data for all analysed Golgi ROIs is available in Figure 9B - Source Data 1. (C) YFP/CFP measurements of GalT-pHlemon in the Golgi ribbon, endo-/lysosomes (cytosolic CFP puncta), as well as in the enlarged endosomes post 20 min nigericin treatment and 100 min recovery. Numerical data for all analysed organelle ROIs is available in Figure 9C - Source Data 1. (D) HeLa cells, stably expressing mApple-Rab5 and transiently transfected with GalT-pHlemon, were treated for 20 min with nigericin, washed and imaged over 3 h. Images show GalT-pHlemon sensor localising to the enlarged transiently Rab5-positive endosome and changing YFP and CFP intensity consistent with endosomal acidification.

### Endosomal acidification is most pronounced during Rab conversion

Equipped with a sensor locating to endosomes and responding to endosomal pH changes, we investigated the kinetics of endosomal acidification relative to Rab5 and Rab7 recruitment, using cells stably expressing mApple-Rab5 or mApple-Rab7 and transiently transfected with GalT-pHlemon. The nigericin-induced enlarged endosomes showed a dramatic decrease in YFP signal, which always coincided with Rab5-positive stages of endosome maturation (Fig 10A, video Fig 10A supplement 1) and with early phases of Rab7 recruitment (Fig 10D, video Fig 10D supplement 1). Ratiometric quantifications of intraluminal YFP and CFP signals revealed relatively stable YFP/CFP ratio prior to Rab5 recruitment, followed by a sharp decrease during Rab5 recruitment and Rab conversion, and stabilisation of a new, lower YFP/CFP ratio in Rab7-positive endosomes (Fig 10B, C, E and F). Conversion of YFP/CFP ratios to pH values indicates an average pH of 6.6 in early endosomes prior to Rab5 recruitment and a final pH of 5.7 in Rab7 endosomes. Our data indicate that the biggest pH drop occurs concomitantly with Rab5-to-Rab7 exchange, pointing to a regulation of acidification during endosome maturation.

**Figure 10.**
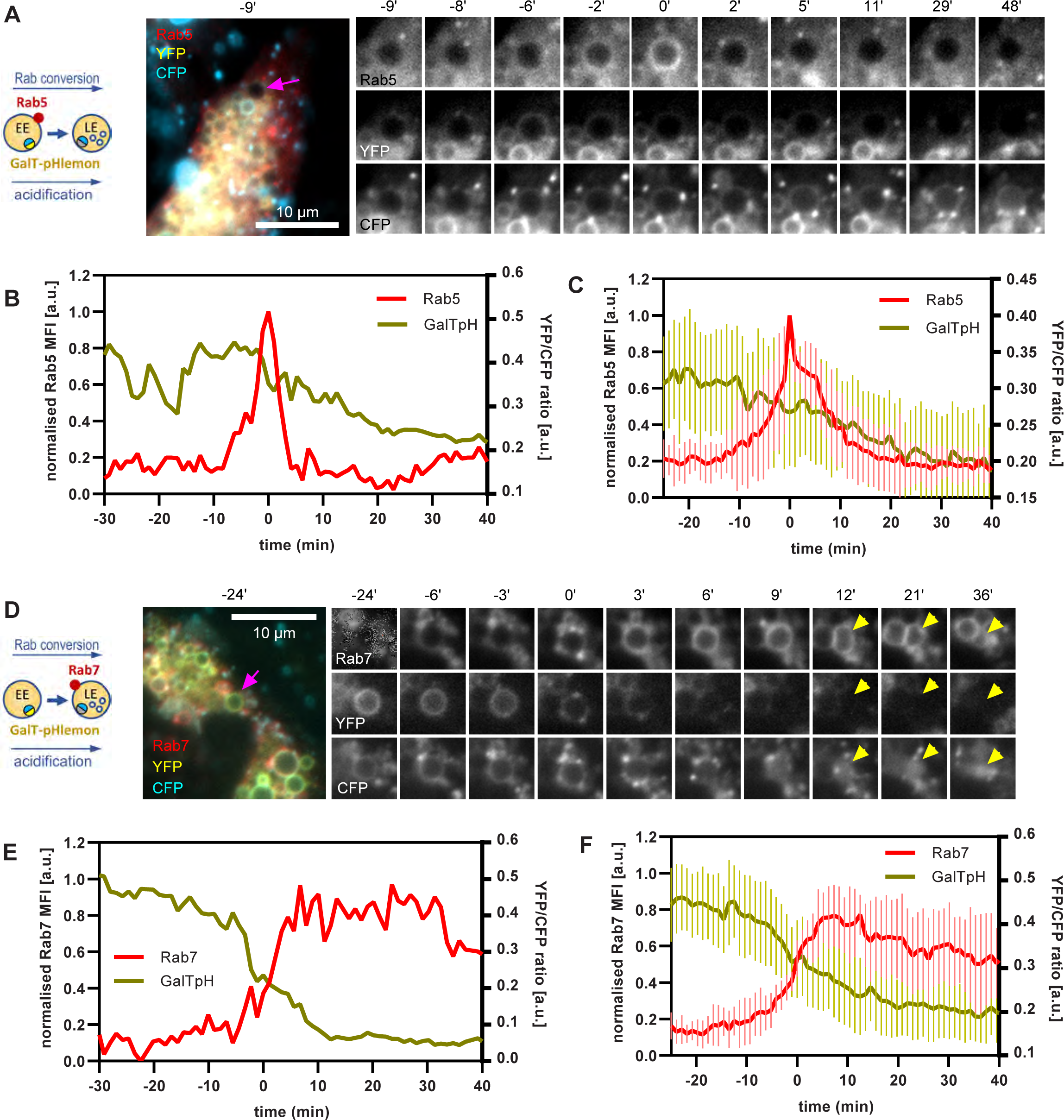
GalT-pHlemon sensor detects endosomal acidification, which correlates with Rab conversion. HeLa cells, stably expressing mApple-Rab5 (A-C) or mApple-Rab7 (D-F) and transiently transfected with GalT-pHlemon, were treated for 20 min with nigericin, washed and imaged over 3 h, as described in Figure 4A. (A,D) Time-lapse images of a representative endosome to show association of Rab conversion with acidification, as detected by the decrease in the YFP signal and relatively constant CFP at the rim. (B,E) Corresponding graphs of normalised mean fluorescence intensity of Rab5 (B) or Rab7 (E) and YFP/CFP ratio of the endosomal GalT-pHlemon signal at the rim of the endosome in (A) or (D), respectively, during and around the time of Rab conversion. (C,F) Averaged Rab5 (C) or Rab7 (F) and GalT-pH kinetics of 15 and 18 endosomes, respectively. Error bars represent standard deviation. Pooled data from two independent experiments. Numerical data for all analysed endosomes is available in Figure 10C - Source Data 1 and Figure 10F - Source data 1.

### Impaired Rab conversion is associated with slower endosomal acidification

If endosomal acidification is dependent on the progression of endosome maturation, then blocking endosome maturation by impairing Rab conversion should undermine acidification. To block Rab conversion, we knocked out the Ccz1, a subunit of the Rab7GEF, which has been shown to promote Rab conversion (Fig. 11, figure supplement 1A) (Nordmann et al., 2010; Poteryaev et al., 2010; van den Boomen et al., 2020). Ccz1 depletion abolished Rab7 recruitment to, and Rab5 removal from, the nigericin-induced enlarged endosomes (Fig 11A, videos Fig 11A supplement 1-3). Ccz1-deficient Rab5-positive endosomes could engage in homotypic fusion and interact with Rab5-positive enlarged compartments but did not mature to classical endolysosomes Figure 11, figure supplement 1B, Fig 1C). These perturbations could be efficiently rescued by expression of wild-type Ccz1 in Ccz1 knock-out cell lines (Fig. 11A, videos Fig 11A supplement 1-3). To ensure that in rescue experiments we selected for analysis only the cells expressing Ccz1, and not untransfected cells, we appended a far-red fluorophore mNeptune2 via the T2A peptide linker to Ccz1, resulting in expression of the two separate proteins in the transfected cells (Fig. 11, figure supplement 2A). The mNeptune2 was tagged with NLS, targeting it to the nucleus, to minimise interference with the mApple signal at the endosomes (Fig. 11, figure supplement 2B). Hence, we have generated Ccz1 knock-out cell lines that showed impaired Rab conversion and could be efficiently rescued with the Ccz1 rescue construct.

**Figure 11.**
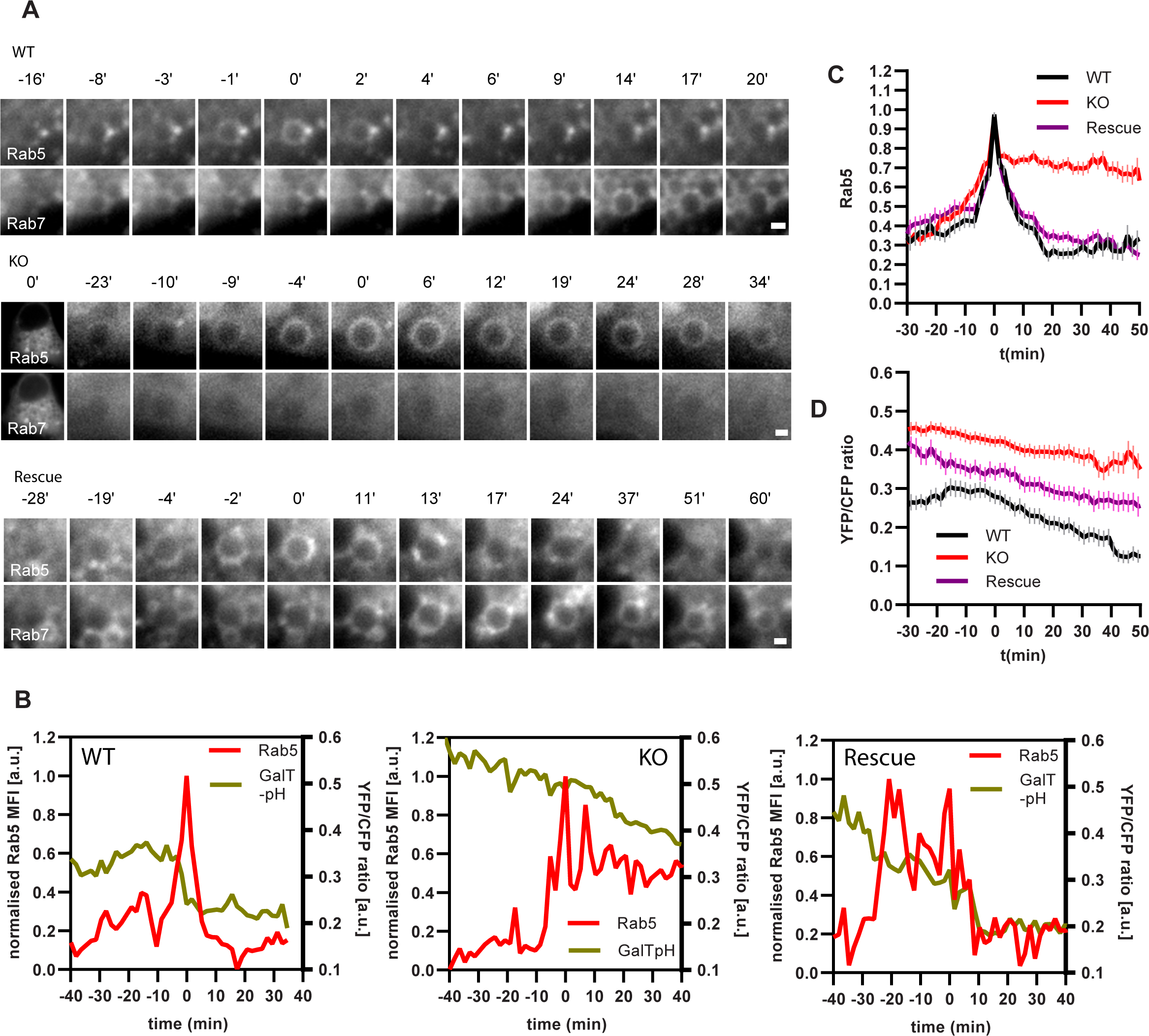
Ccz1 KO disrupts Rab conversion and delays endosomal acidification. HeLa cell lines with wild-type (WT) Ccz1 and knocked-out Ccz1 (KO) were transiently transfected with mApple-Rab5 and either GFP-Rab7 (A) or GalT-pHlemon (B-D). Ccz1 expression plasmid was co-transfected for 72 h for rescue experiments. Nigericin was added to cells for 20 min and washed away, and cells were imaged by time-lapse microscopy, as described in Figure 4A. (A) Time-lapse images of representative endosomes to show absence of Rab7 recruitment and the associated lack of displacement of Rab5 in KO cells, compared to the expected Rab conversion in WT and rescue cells. Scale bar = 1 µm. (B) Graphs of normalised mean fluorescence intensity of Rab5 and YFP/CFP ratio of the endosomal GalT-pHlemon signal in representative endosomes during and around the time of Rab conversion / Rab5 peak, in WT, KO and rescue cells. (C,D) Averaged kinetics of Rab5 recruitment (C) and corresponding GalT-pHlemon YFP/CFP ratios (D) for WT, KO and rescue cells, in 54, 54 and 56 endosomes, respectively. Error bars represent SEM. Pooled data from three independent experiments using different Ccz1 clones. Numerical data for all analysed endosomes is available in Figure 11 - Source Data 1.

To test our hypothesis that impaired Rab conversion compromises endosomal acidification, we expressed mApple-Rab5 and GalT-pHlemon in Ccz1-KO and control cells, and monitored the YFP/CFP ratio kinetics of the pH sensor on Rab5-positive endosomes (Fig 11B, Fig. 11, figure supplement 1B). While in the control and Ccz1 rescue cells, acidification occurred with similar kinetics as observed above (Fig 10B and C), in Ccz1 KO cells the acidification was strongly delayed (Fig 11B-D). Nevertheless, acidification occurred eventually after a long delay in Ccz1 KO cells. In line with this conclusion, Ccz1 KO cells have grossly enlarged CFP-filled puncta and compartments, reflective of their enlarged lysosomes and acidified hybrid compartments as also observed with Lysotracker staining of Rab5-positive compartments (Fig 12A, Fig 12, figure supplement 1). The CFP-positive compartments in untreated cells showed no differences in pH between Ccz1-replete and Ccz1-deficient cells, ranging from pH 5.7 to pH 4.0, indicative of endolysosomes and lysosomes, respectively (Fig 12B). Furthermore, following disruption of pH by nigericin treatment and washout, wild-type cells restored their lysosomal pH, while Ccz1 KO cells displayed a wide range of pH in pHlemon-filled compartments, ranging from pH 6.4 to pH 4.0 (Fig 12C and D). Taken together, our data suggest that Rab conversion is driving efficient endosomal acidification.

**Figure 12.**
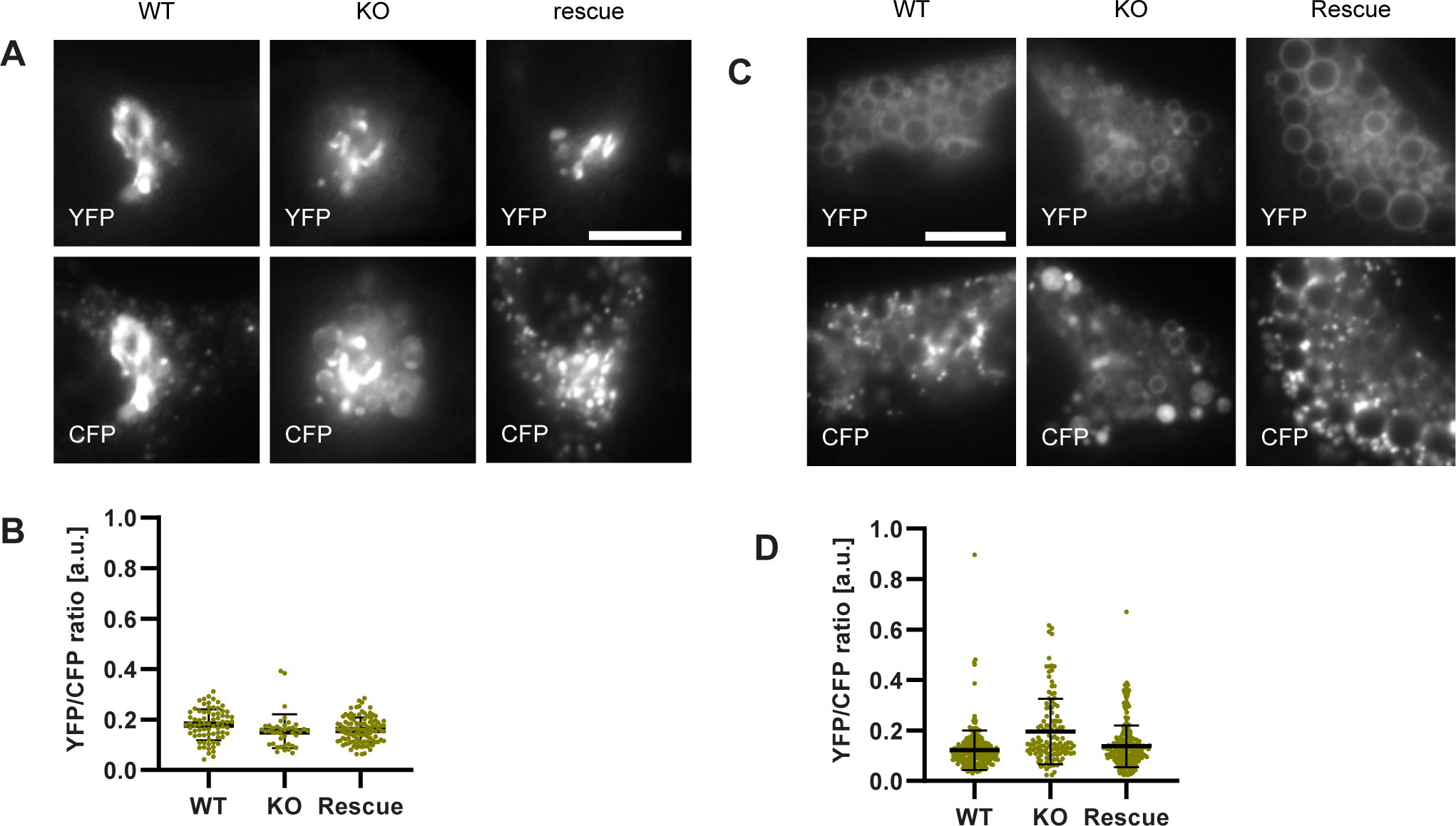
Lysosomes of Ccz1 knockout cells can acidify to the same extent as wild-type cells, with some delay. HeLa cell lines with wild-type (WT) Ccz1 and knocked-out Ccz1 (KO) were transiently transfected with GalT-pHlemon. Ccz1 expression plasmid was co-transfected for 72 h for rescue experiments. WT, KO and rescue cells were left untreated (A,B) or treated with 20 min nigericin followed by 50 min recovery (C,D). (A) Images of cells showing Golgi and highly acidified organelles as visualised with GalT-pHlemon. Acidified organelles appear as puncta in WT and rescue cells and as larger round CFP-filled compartments in KO cells. Scale bar = 10 µm. (C) Images of cells pre-treated with nigericin have dispersed trans-Golgi and re-acidified organelles as in (E). Scale bar = 10 µm. (B,D) Corresponding measurements of YFP/CFP ratio of the GalT-pHlemon sensor in the acidified organelles. Numerical data for all analysed endosomes is available in Figure 12B - Source Data 1 and Figure 12D - Source Data 1.

## DISCUSSION

Despite the pivotal importance of endosomal transport for cell survival and communication, tissue organization and development in all eukaryotic cells, endosomes largely escape largely precise temporal and spatial observations because of their small size and mobility. Several approaches using enlarged endosomes with low motility have successfully been used in the past, however, these were restricted to a particular cell type and organism or a limited subset of endosomes (Poteryaev et al., 2010; Skjeldal et al., 2021; van Weering et al., 2012). We report here an easy, reliable and inexpensive method to enlarge endosomes in a variety of different cell types and follow them over their entire lifetime - from initial formation and subsequent maturation to endolysosome formation and lysosome maturation. Importantly, our acute nigericin treatment takes advantage of the natural, physiological cellular stress response and does not cause any autophagy induction, lysosomal damage, cell death or growth retardation. Moreover, this assay does not require any special equipment; a standard fluorescence microscope equipped with commonly used filters and a camera are sufficient. We validated our assay by comparing our measured parameters of endosome maturation with published data. Most importantly, Rab5-to-Rab7 conversion progressed with previously published kinetics (Poteryaev et al., 2010; Rink et al., 2005; Skjeldal et al., 2021; van der Schaar et al., 2008). In addition, PI(3)P was produced on Rab5 endosomes at about the same rate, in which Rab5 levels increased, consistent with the positive feedback loop between Rab5 and the PI(3)P kinase Vps34 (Zerial and McBride, 2001). While the PI(3)P levels coincided with the Rab5 peak, we observed a short delay in decrease of the FYVE-GFP. This finding is consistent with a previous report, which described *C. elegans* Rab7GEF recruitment to PI(3)P-rich endosomes (Poteryaev et al., 2010) to drive Rab conversion. Additionally, Vps34 removal and subsequent PI(3)P turnover is dependent on low luminal pH as characteristic of late endosomes (Naufer et al., 2018; Podinovskaia et al., 2013). It is conceivable that the slight delay in reducing PI(3)P levels might provide directionality during Rab conversion. Thus, our assay faithfully recapitulates endosome maturation as described in other model systems.

We used the established assay to determine whether there is a strict coordination between recycling to the plasma membrane and Rab conversion. There is evidence in the literature that recycling to the plasma membrane only happens from Rab5 compartments or from Rab5 and Rab7 compartments (van Weering et al., 2012). Our data indicate that recycling to the plasma membrane can happen throughout the lifetime of an endosome. We observed that SNX1 was recruited simultaneously with Rab5 probably mediated through its PX domain (Peter et al., 2004), consistent with the reported coordination between the two proteins (van Weering et al., 2012). The PX domain recognises PI(3)P, which is recruited concomitantly with Rab5 to maturing endosomes. However, even though in about one third of the endosomes, there seemed to be temporal coordination between SNX1 and Rab5 removal from the endosome, sorting persisted in the remaining Rab7-positive endosomes. In addition, we did not observe any spatial coordination on the endosomal membrane as the SNX1 and Rab5 or Rab7 domains appeared to move independently. Moreover, Rab11 contacted maturing endosomes irrespective of whether they were Rab5 or Rab7 positive. Therefore, our data suggest that the onset of recycling is coordinated with the arrival of Rab5, at least for Snx1, but the process itself is independent of Rab conversion, as previously suggested (Rojas et al., 2008). Consistently, we have shown previously that when Rab conversion is blocked, Rab11 localization and Rab11-dependent recycling are not affected in *C. elegans* oocytes (Poteryaev et al., 2010; Poteryaev et al., 2007).

Although acidification of endosomes is required for endosome maturation (Podinovskaia and Spang, 2018), how this process is regulated remains poorly understood. Since Rab conversion is a major driver of endosome maturation, we asked whether Rab conversion regulates endosomal acidification. Unfortunately, all pH sensor probes we tried turned out not to be useful because they were mostly stuck in the ER. While in neurons, in which the probes have mainly been applied, this might be less of an issue, in our system this has prevented any meaningful analysis. We, hence, developed a new probe based on the ratiometric pHlemon and GalT, which localises to Golgi but enters also the endolysosomal pathway. With this new probe, we showed that Rab conversion is required for efficient acidification. Over extended times, acidification of endosomes still occurred in absence of Rab conversion and we speculate that this acidification can help drive fusion with lysosomes. Moreover, this effect may explain why a *sand-1* mutant in *C.elegans*, knockdown of Mon1a and b, or a Ccz1 KO in mammalian cells is not lethal (Poteryaev et al., 2010; Poteryaev et al., 2007). How Rab conversion promotes a drop in pH, we can only speculate at this point. It is possible that the activity of the V-ATPase is upregulated during Rab conversion, driven by Rab7 effectors such as RILP (Bucci et al., 2000; De Luca and Bucci, 2014; De Luca et al., 2014), and is accompanied by the arrest in interactions with less acidified endocytic vesicles. It is also conceivable that there is a simple regulatory loop such as phosphorylation and dephosphorylation of V-ATPase subunits, however, the potential kinase or phosphatase regulators remain to be identified. Additionally, proton channels such as Nhe6 have been reported to finetune endosomal pH, and their role in endosome acidification kinetics remains to be explored. Finally, multiple factors are known to interact with the V-ATPase; their role in controlling the acidification during Rab conversion remains to be established.

In conclusion, we have developed and validated a straightforward live-cell imaging assay and used it to define kinetics and regulation of mediators of endosomal maturation. This assay will be invaluable for addressing outstanding questions relating to the regulation and potential coordination of processes during endosome maturation. Given its applicability to different cell types, it may also be useful in establishing cell-type specific differences in the regulation of endosomal transport. The assay has been designed to make it accessible and applicable in most cell biology laboratories and proved to be a powerful tool to further our understanding of endosome biology.

## MATERIALS AND METHODS

### Cell culture

HeLa CCL2 and HeLa Kyoto-α cells were grown at 37°C and 5% CO_2_ in high-glucose Dulbecco’s modified Eagle’s medium (DMEM, Sigma-Aldrich) supplemented with 10% fetal calf serum (FCS, Biowest), 2 mM L-Glutamine, 1 mM Sodium Pyruvate, and 1x Penicillin and Streptomycin (all from Sigma).

For transient cell transfections, cells were plated into 6-well plates to reach 70% confluency the following day and transfected with 0.5 µg plasmid DNA complexed with Helix-IN transfection reagent (OZ Biosciences). Cells were used in imaging assays at 48 h post transfection. For the Ccz1 rescue experiments, given the larger size of the plasmid, 2 µg plasmid DNA was used, and cells were transfected for 72h.

For cell growth assays, cells were plated into 12-well plates at 10,000 cells per well. The following day, a sample was taken for counting (0 h time point), and the remaining cells were incubated in complete medium with or without nigericin for 20 min, washed, incubated for specified time periods and collected for counting. Both conditions were reaching 90% confluency by 72 h. Three wells were measured per condition. Doubling time was calculating using the formula

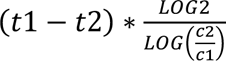

where c1 and c2 are cell counts for two consecutive time points t1 and t2.

### DNA constructs

mApple-Rab5a, mApple-Rab7a, GalT-mCherry, mNeptune2-C1 and mApple-Lamp1-pHluorin plasmids were a gift from Michael Davidson (Addgene # 55944, 54945, 55052, 54836, 54918). GFP-Rab11a was a gift from Richard Pagano (Addgene # 12674) (Choudhury et al., 2002). EGFP-2xFYVE was a gift from Harald Stenmark (Addgene # 140047) (Gillooly et al., 2000). Lamp1-GFP was a gift from Esteban Dell’Angelica (Addgene #34831) (Falcon-Perez et al., 2005). pSpCas9(BB)-2A-GFP (pX458) and pSpCas9(BB)-2A-puro (pX459) were a gift from Feng Zhang (Addgene #48138 and #48139) (Ran et al., 2013). Snx1-turboGFP and Ccz1-myc were from Origene (#RG201844 and RC222195). GFP-Rab7a was generated by substituting mApple in the Rab7a plasmid with GFP from GFP-Rab11a using NheI and XhoI restriction sites.

For stable GalT-GFP cell line generation, a sequence encoding GalT-EGFP (B4GALT1 ORF minus the catalytic moiety) was generously provided by Jennifer Lippincott-Schwartz (Howard Hughes Medical Institute, Ashburn, VA). GalT-EGFP was amplified using primers with restriction site overhangs for NotI and PacI, and subcloned into the Retro-X Q vector pQCXIP (Takara Bio). The GalT-EGFP plasmid for transient transfections was generated from GalT-mCherry plasmid and the EGFP insert from LAMP1-EGFP, using NEBuilder HiFi Assembly cloning kit (New England Biolabs, NEB) with the primers designed by the NEBuilder Assembly Tool (table 1) following manufacturer’s instructions.

**table 1.**
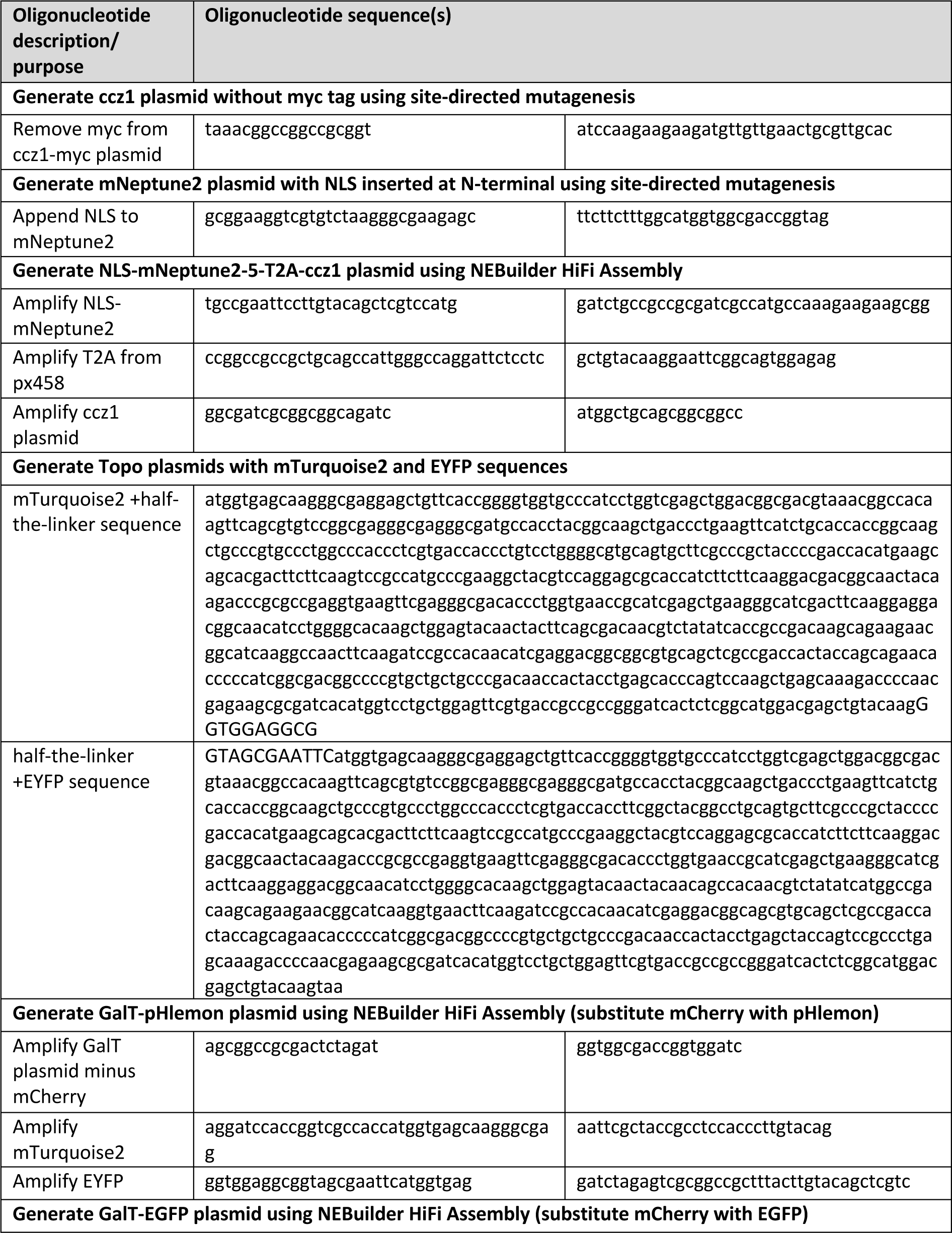
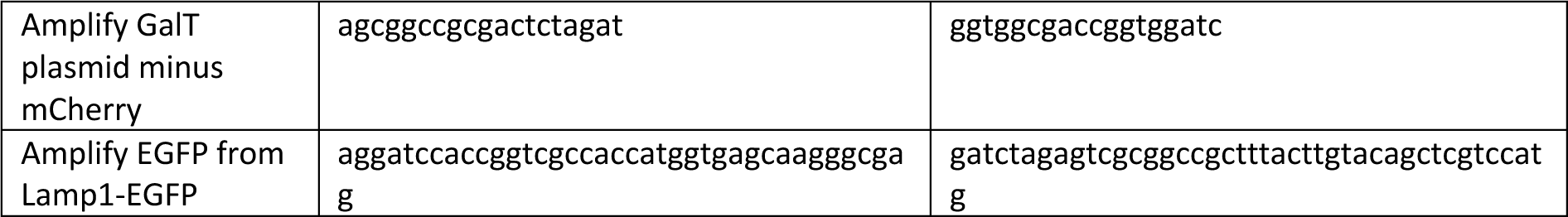
Oligonucleotide sequences used to generate specified constructs

For Ccz1 rescue experiments, the myc tag was removed from the Ccz1-myc plasmid using NEB site-directed mutagenesis (SDM) kit (NEB) following manufacturer’s instructions and primers selected using NEBaseChanger tool (table 1). Nuclear localisation sequence (NLS) was cloned at the N terminal of mNeptune2 using NEB SDM kit with the mNeptune2-C1 plasmid following manufacturer’s instructions and primers listed in table S1. To generate NLS-mNeptune2-T2A-ccz1 and NLS-mNeptune2-T2A-ccz1-myc, primers were designed with NEBuilder Assembly Tool (NEB) as listed in table 1 to generate PCR products from NLS-mNeptune2, pSpCas9(BB)-2A-GFP and ccz1 plasmids. Purified PCR products for NLS-mNeptune2 and T2A peptide were ligated together using forward primer for mNeptune2 and reverse primer for T2A in a PCR reaction with 5 cycles at 50°C and 25 cycles at 63.7°C (i.e. the annealing temperature of the two primers). Purified NLS-mNeptune2-T2A and ccz1 backbone PCR products were assembled together using NEBuilder HiFi Assembly cloning kit (NEB).

For the pH sensor constructs, we used pHlemon, which consists of mTurquoise2 and EYFP in tandem with a 21bp linker in between (Burgstaller et al 2019). Separate constructs for mTurquoise2+half-the-linker and half-the-linker+EYFP were synthesized by Twist Bioscience (table 1). The two sequences were cloned separately into pCR Blunt II-Topo vectors (Invitrogen). For the GalT-pHlemon plasmid, primers were designed with NEBuilder Assembly Tool (table 1) to generate PCR products for the mTurquoise2 and EYFP as well as the GalT backbone without the tag from GalT-mCherry plasmid. Purified PCR products were assembled together using NEBuilder HiFi Assembly cloning kit.

### Fluorescent cell line generation

To generate GalT-EGFP stable cell lines, Phoenix Ampho packaging cells (from the Nolan lab, Stanford University) were grown in complete medium supplemented with 1 mM sodium pyruvate and transfected with pQCXIP-GalT-EGFP using FuGENE HD (Promega) to produce viral particles. The viral supernatant was harvested after 48-72 h, passed through a 0.45 µm filter, supplemented with 15 µg/ml polybrene, and added to target HeLa-α cells. The next day, complete medium with 1.5 µg/ml puromycin was added. Following selection, cells were subjected to cell sorting on a FACSAria III (BD Biosciences) to obtain a cell pool with homogenous expression levels. HeLa-*α*-GalT-EGFP were maintained in complete medium supplemented with 1.5 µg/ml puromycin at 37°C in 5% CO_2_.

Stable expression of mApple-Rab5 in HeLa cells was achieved by transfection of HeLa CCL2 cells with the mApple-Rab5a plasmid and three rounds of bulk-sorting by FACS at 15, 33 and 61 days post transfection. The cell line stably expressing mApple-Rab7a were generated as previously described (Wu et al., 2020). Briefly, HeLa CCL2 cells were transfected with the mApple-Rab7a plasmid, FACS-sorted for mApple-positive cells at 20 days and into 96-well plates 15 days later for clonal expansion. To generate the cell line with stable expression of both, mApple-Rab5 and GFP-Rab7, stably-expressing mApple-Rab5 cells were transfected with GFP-Rab7 and FACS-sorted at 7 days post transfection for a bulk population of mApple-positive and GFP-positive cells and, a further 11 days later, into 96-well plates for clonal expansion. The doubly-positive colonies were bulk-sorted once again 50 days later to remove cells that were no longer expressing either marker.

### CRISPR-Cas9 knock-out of Ccz1

Ccz1 has a homolog, Ccz1b, which differs by 4 nucleotides and is identical in amino acid sequence. Guide RNAs (gRNA) were designed to target both genes simultaneously. The CRISPR strategy was described previously (Ran et al., 2013; Solinger et al., 2020). Briefly, based on gRNAs targeting the first and last exons of Ccz1, two double-stranded oligonucleotide sequences were cloned one each into the two plasmids, Px458 (GFP) and Px459 (Puro) and transfected into HeLa CCL2 cells. Plasmids without the inserts were used as controls. After 24 h of transfection, cells underwent 24 h selection with puromycin, followed by single-cell FACS sorting of GFP-positive cells for clonal expansion. Clones showing >90% reduction in Ccz1 expression were used for evaluation of endosome maturation. Three different ccz1-deficient (KO) clones and three different control clones (WT) were used in experiments to obtain the three biological replicates.

### Live cell imaging

Cells were plated into 8-well imaging chambers (Miltenyi) the day before imaging to reach 50-70% confluency on the day of imaging. Just prior to imaging, cell medium was replaced with pre-warmed imaging buffer (5 mM Dextrose, 1 mM CaCl_2_, 2.7 mM KCl, 0.5 mM MgCl_2_ in PBS supplemented with 10% FCS and Penicillin and Streptomycin). Where specified, cells were treated with 10 µM Nigericin (AdipoGen, 10 mM stock in Ethanol) or 5 µM Monensin (Sigma, 50 mM stock in Ethanol) prior or during imaging. Where specified, 100 nM Lysotracker Green (Molecular Probes, 1mM stock in DMSO) or 10nM Lysotracker Red in imaging buffer was added 20 min prior to imaging.

Cells were imaged at 37 °C using an inverted Axio Observer microscope (Zeiss) with a Plan Apochromat N 63×/1.40 oil DIC M27 objective and a Photometrics Prime 95B camera. Filters with standard specifications for CFP, GFP, YFP, dsRed and APC were used to image mTurquoise2, GFP, YFP, mApple and mNeptune2, respectively. To minimise overexpression artifacts, cells were selected for imaging that expressed minimal amount of each fluorescent marker that was sufficient to produce good quality signal. For time-lapse experiments, to monitor endosome maturation kinetics, cells were treated with nigericin for 20 min, washed 4 times in imaging buffer and imaging chamber mounted onto an automated microscope stage. Several fields of view (FOV) were selected for imaging per condition. The microscope was programmed to image all FOVs sequentially and repeat the imaging at 1-2 min intervals. The obtained images corresponded to the recovery time between 40 min and 190 min post nigericin treatment. For experiments comparing Ccz1 WT to KO clones, with and without rescue, all four conditions were imaged in a single time-lapse experiment. All experiments were performed three times on different days.

### Single endosome analysis and quantification

Using the time-lapse images, endosomes were selected for analysis with the following criteria. For mApple-Rab5 expressing cells, endosomes initially devoid of Rab5 and later acquiring and subsequently losing Rab5 were identified. For mApple-Rab7 expressing cells, endosomes initially devoid of Rab7 and subsequently acquiring Rab7 and stabilising its expression were identified. This ensured that the entire Rab conversion event was captured in the kinetic.

To quantify the recruitment of markers to the endosome, an oval selection tool in Fiji was used to draw a circular region-of-interest (ROI) closely following the rim of the endosome in the channel for the most visible marker or by predicting the location of the rim if cases where the enlarged endosome was negative for both markers and appeared as a dark circle (Fig 4A). For less circular endosomes and endolysosomes, the ROI was adjusted using the elliptical or a free-hand selection tool. ROIs were adjusted for every time point where the rim of the endosome could be unambiguously identified. Mean fluorescence intensity (MFI) of a two-pixel wide rim at the ROI was recorded in all channels. A larger two-pixel wide rim three pixels away from the endosome was generated with a macro based on the original ROI, and MFI was calculated as a measure of background for each time point (Fig 4A). We found that adjusting the MFI at the rim of endosome for this background MFI minimised noise and produced data reflective of visual assessment of marker presence at the endosome. For intraluminal pHlemon measurements, the circular ROI at the rim of the endosome was reduced by 1 pixel and the total MFI of the reduced ROI was calculated in both YFP and CFP channels. A ROI in a field where no cells were present was measured to obtain background values. This approach was found to produce pH measurements as accurate as the modified rim measurements, in which select pixels were removed to exclude interference from the highly acidified vesicles interacting with the enlarged endosome. For the subdomain measurements, two-pixel thick segmented lines with spline fit were drawn around the full perimeter of the endosome starting at the top, and histogram measurements were obtained of fluorescence intensity along the length of the line.

Endosomal recruitment marker measurements were background-subtracted and adjusted for the minimum and maximum values of the entire measured kinetic, to represent a range from 0 to 1. The pHlemon measurements were kept as background-adjusted YFP/CFP ratios. For averaging, kinetics were aligned for Rab5 peak or for Rab7 at 50% of its final maximum value, representing the point of Rab conversion (Fig 4C-D). At least 10-20 endosomes from at least 3-10 cells were used in analysis and each experiment was repeated at least 3 times. Means, standard deviations, SEM, Pearson’s correlation and linear regression were calculated in GraphPad Prism.

### pH calibrations

GalT-pHlemon images provided us with pH-responsive YFP/CFP ratio measurements. To convert YFP/CFP ratios to pH values, GalT-pHlemon expressing cells were incubated in buffers of known pH containing 5 µM Nigericin, 50 µM Monensin and 100 nM Concanamycin A (Sigma, 100 µM stock in DMSO) to disrupt intracellular proton gradients. The buffers consisted of 138 mM NaCl, 5 mM KCl, 2 mM CaCl_2_, 2 mM MgCl_2_, 10 mM Glucose, 10 mM HEPES (for pH 5.5-9.0) or 10 mM MES (for pH 4.0-5.0), and pH was adjusted with NaOH or HCl. Images were taken at 15 min after adding the buffers to the cells. Golgi ribbon structures were selected with a segmented line tool in Fiji, and MFI was calculated in both channels and subtracted for background measured in a field devoid of cells. The equation for the line of best fit was generated in GraphPad Prism based on log(dose) response curve with variable slope, where log(dose) is pH and response is YFP/CFP values (sigmoidal four-parameter dose-response curve; Fig S9).

### Electron microscopy

HeLa CCL2 cells were grown in 10 cm dishes, treated for 20 min with nigericin and left to recover. At specified times, cells were fixed in DMEM containing 2.5% glutaraldehyde and 3% formaldehyde for 2 h at room temperature. Cells were washed with PBS and cell membranes stained with 2% osmium tetroxide and 1% potassium hexacyanoferrate in H_2_O for 1 h at 4°C. Following a wash with water, cells were further stained for proteins and nucleic acids with 2% uranyl acetate in H_2_O overnight at 4°C. Samples were subsequently dehydrated in acetone/H_2_O in stepwise increases in acetone concentration of 20%, 50%, 70%, 90% and 3x 100%. The samples were infiltrated with 50% epon embedding medium in acetone for 1 h at room temperature, and subsequently with 100% epon resin overnight. Next day, fresh epon resin was added and samples were polymerised for 24 h at 60°C. Sections of 60–70 nm were collected on carbon-coated Formvar-Ni-grids and were viewed with a Phillips CM100 electron microscope

To prepare cells for immunolabeling, HeLa cells stably expressing GalT-GFP were prepared as previously described (Beuret et al., 2017). Sections were stained sequentially with rabbit anti-GFP (1:100; Abcam 6556) and goat anti-rabbit coupled to 10 nm gold particles (BB International). For dual labelling, HeLa cells stably expressing mApple-Rab5 were transiently transfected with GalT-EGFP, prepared for immunolabelling as above, and stained sequentially for GalT-GFP and mApple-Rab5. The sections were blocked with PBST (PBS+0,05% Tween20) supplemented with 2% BSA for 20 min, incubated overnight at 4°C with anti-GFP (1:100, Abcam), washed five times for 5 min with PBS and incubated for 2 h at room temperature (RT) with donkey anti-rabbit coupled with 12 nm Gold (Jackson Immuno Research). The wash step was repeated and the single-stained sections were subsequently fixed for 2 min with 1% glutaraldehyde in PBS, washed with PBS and blocked for 10 min with the blocking solution. Following a 2 h incubation at RT with anti-mCherry (1:100, St John’s Laboratory STJ140001), the sections were washed with PBS, blocked with PBS supplemented with 2% fish gelatin (Sigma) for 10 min and incubated with mouse anti-goat (1:100, Jackson Immuno Research) for 90 min. The wash and the 2% fish gelatin block steps were repeated and the sections were incubated with goat anti-mouse coupled with 5 nm Gold (1:100, BBInternational) for 2 h. The double-stained sections were washed five times for 5 min with PBS and three times for 2 min with H_2_O, and subsequently stained for 10 min with 2% uracyl acetate and 2 min Reynold’s solution.

### Immunostaining

HeLa cells were plated onto coverslips 24 h prior to 20 min nigericin treatment and recovery. At specified times post nigericin treatment, cells were fixed in 2% paraformaldehyde, permeabilised with 0.1% Triton X-100, blocked in PBS containing 2% BSA and 5% goat serum, and stained with anti-GalT (1:50, Sigma HPA010807) followed by AF594-conjugated goat anti-rabbit and DAPI. Coverslips were mounted onto glass slides with Fluoromount G (Southern Biotech) and sealed with nail polish. Following z-stack image acquisition in DAPI and dsRed channels, images were deconvolved using Huygens deconvolution online software tool.

## Supplementary methods

### Cell culture

HEK293 and Neuro2A cells were grown at 37°C and 5% CO2 in high-glucose Dulbecco’s modified Eagle’s medium (DMEM, Sigma-Aldrich) supplemented with 10% fetal calf serum (FCS, Biowest), 2 mM L-Glutamine, 1 mM Sodium Pyruvate, and 1x Penicillin and Streptomycin (all from Sigma). Cos-1 cells were grown in low-glucose DMEM supplemented as above. For live cell microscopy, imaging chambers were pre-coated with poly-L-lysine to enhance attachment of these three cell types.

### DNA constructs

mTagBFP2-Rab7 was generated by replacing Rab5 in mTagBFP2-Rab5a (Addgene #55322, gift of Michael Davidson) with Rab7 from mApple-Rab7 (Addgene #54945) using BamHI and XhoI restriction sites.

### Live cell imaging

For endolysosomal labelling, 0.5 mg/mL Dextran-AF488 (10,000 MW, Molecular Probes, 10 mg/mL stock in water) was pulsed into cells for 17 h and chased for 4 h into lysosomes. For endosomal dextran labelling, 0.5 mg/mL Dextran-AF488 was pulsed into cells for 65 min post nigericin treatment and subsequently washed away for imaging within the following 15 min.

### qRT-PCR

RNA was extracted and purified from cells using RNeasy kit or using the Trizol-Chloroform extraction and isopropanol precipitation, following manufacturer’s instructions. cDNA was reverse-transcribed using GoScript reverse transcriptase primed with a mix of Oligo(dT)s and random hexamers (Promega). qRT-PCR was performed using GoTaq qPCR master mix (Promega) and primers specific for spanning the intron junction between exons 3 and 4 of ccz1 (ACATTTAGCCCATCAAAACCTGC, CCGAACAACCATGACCATCC). Ccz1 expression was normalized for β-actin expression.

### Western blotting and antibodies

HeLa cells transiently transfected with NLS-mNeptune2-5-T2A-ccz1-myc were lysed in lysis buffer (1% NP-40, 150 mM NaCl, 50 mM Tris pH8.0, protease inhibitors) and denatured in Laemmli buffer at 65°C for 5 min. Samples were resolved by SDS-PAGE and transferred onto nitrocellulose membrane. Samples were blocked in milk, stained with anti-myc (1:2,000, Sigma 9E10) and HRP-conjugated anti-mouse, and revealed with WesternBright ECL HRP substrate (Advansta).

### Immunofluorescence

HeLa cells stably expressing mApple-Rab5 were plated onto coverslips 24 h prior to treatment. Cells were left untreated, treated for 16 h with bafilomycin (100 nM,Enzo Life Sciences, 100 μM stock in DMSO), or treated with nigericin for 20min followed by a 60 min washout. Following treatment, cells were fixed for 10 min with methanol at -201C, blocked in PBS containing 2% BSA and 5% goat serum, and stained with anti-LC3b (1:400, Cell Signaling Technology #3868) followed by AF488-conjugated goat anti-rabbit. Coverslips were mounted onto glass slides with Fluoromount G (Southern Biotech) and sealed with nail polish. Following z-stack image acquisition, images were deconvolved using Huygens deconvolution online software tool.

## Acknowledgements

We are grateful for the support provided by the Biozentrum FACS and Imaging core facilities, in particular to Janine Bögli, Stella Stefanova and Laurent Guerard. We thank J. Lippincott-Schwartz for the GalT-EGFP plasmid. This work was supported by the Swiss National Science Foundation (CRSII3_141956 and 310030_197779 to AS) and the University of Basel.

## Supplementary Figures

Figure 1,Figure Supplement 1. Enlarged compartments interact with early endosomes and acquire intraluminal content from the endolysosomal pathway.

Figure 1,Figure Supplement 2. Enlarged endosomes that undergo Rab conversion can be induced by different treatments.

Figure 1,Figure Supplement 3. Enlarged endosomes that undergo Rab conversion can be induced in different cell types.

Figure 2,Figure Supplement 1. Cell growth is unaffected by nigericin treatment, which induces reversible Golgi vesicula1on.

Figure 4,Figure Supplement 1. Enlarged endosomes occasionally display multiple Rab5 peaks and form Rab5 subdomains.

Figure 5,Figure Supplement 1. PI(3)P is recruited to endosomes concomitantly with Rab5.

Figure 6,Figure Supplement 1. Dynamic Snx1 recruitment suggest active sorting at the enlarged endosome.

Figure 8,Figure Supplement 1. Rab11 interacts with the maturing endosome independently of Rab5 or Rab7.

Figure 9,Figure Supplement 1. GalT-pHlemon sensor responds to pH changes in a sigmoidal dose-response manner.

Figure 11, Figure Supplement 1. Validation and characterisation of Ccz1 knockout cell lines.

Figure 11, Figure Supplement 2. Validation and characterisation of Ccz1 the rescue construct.

Figure 12, Figure Supplement 1. Characterisation of Ccz1 knockout cell lines.

## Source data files

Figure 4D - Source Data 1. Quantification of Rab5 and Rab7 recruitment at endosomes.

Figure 5C - Source Data 1. Quantification of Rab5 and PI(3)P recruitment at endosomes.

Figure 6C - Source Data 1. Quantification of Rab5 and Snx1 recruitment at endosomes.

Figure 6E - Source Data 1. Quantification of Rab5 and Snx1 subdomains at endosomes.

Figure 7 - Source Data 1. Quantification of Rab7 and Snx1 recruitment at endosomes.

Figure 8C - Source Data 1. Quantification of Rab5 and Rab11 recruitment at endosomes.

Figure 8D and S8 - Source Data 1. Quantification of Rab5 and Rab11 subdomains at endosomes.

Figure 8G - Source Data 1. Quantification of Rab7 and Rab11 recruitment at endosomes.

Figure 8H and S8 - Source data 1. Quantification of Rab7 and Rab11 subdomains at endosomes.

Figure 9B - Source Data 1. Quantification of GalT-pHlemon signal in cells in calibration buffers of known pH.

Figure 9C - Source Data 1. Quantification of GalT-pHlemon signal in Golgi ribbon, endo-/lysosomal puncta, and enlarged endosomes.

Figure 10C - Source Data 1. Quantification of Rab5 recruitment and GalT-pHlemon signal at endosomes.

Figure 10F - Source data 1. Quantification of Rab7 recruitment and GalT-pHlemon signal at endosomes.

Figure 11 - Source Data 1. Quantification of Rab5 recruitment and GalT-pHlemon signal at endosomes in Ccz1 WT, KO and rescue cells.

Figure 12B - Source Data 1. Quantification of GalT-pHlemon signal in endo-/lysosomal puncta in Ccz1 WT, KO and rescue cells.

Figure 12D - Source Data 1. Quantification of GalT-pHlemon signal in endo-/lysosomal puncta in Ccz1 WT, KO and rescue cells post nigericin treatment.

Figure S4C - Source Data 1. Cell growth numbers and quantification post nigericin treatment.

Figure S7C - Source Data 1. Quantification of Rab5 and Snx1 subdomains at endosomes.

Figure S7E - Source Data 1. Quantification of Rab7 and Snx1 subdomains at endosomes.

Figure S9 - Source Data 1. Sigmoidal dose-response curve to define relationship between GalT-pHlemon signal and pH.

Figure S10A - Source Data 1. Raw RT-PCR data for Ccz1 expression levels in Ccz1 WT vs KO cells.

Figure S11A - Source Data 1. Original Western Blot images for NLS-mNeptune-T2A-myc expression.

---

**Figure 1, Figure supplement 1.**
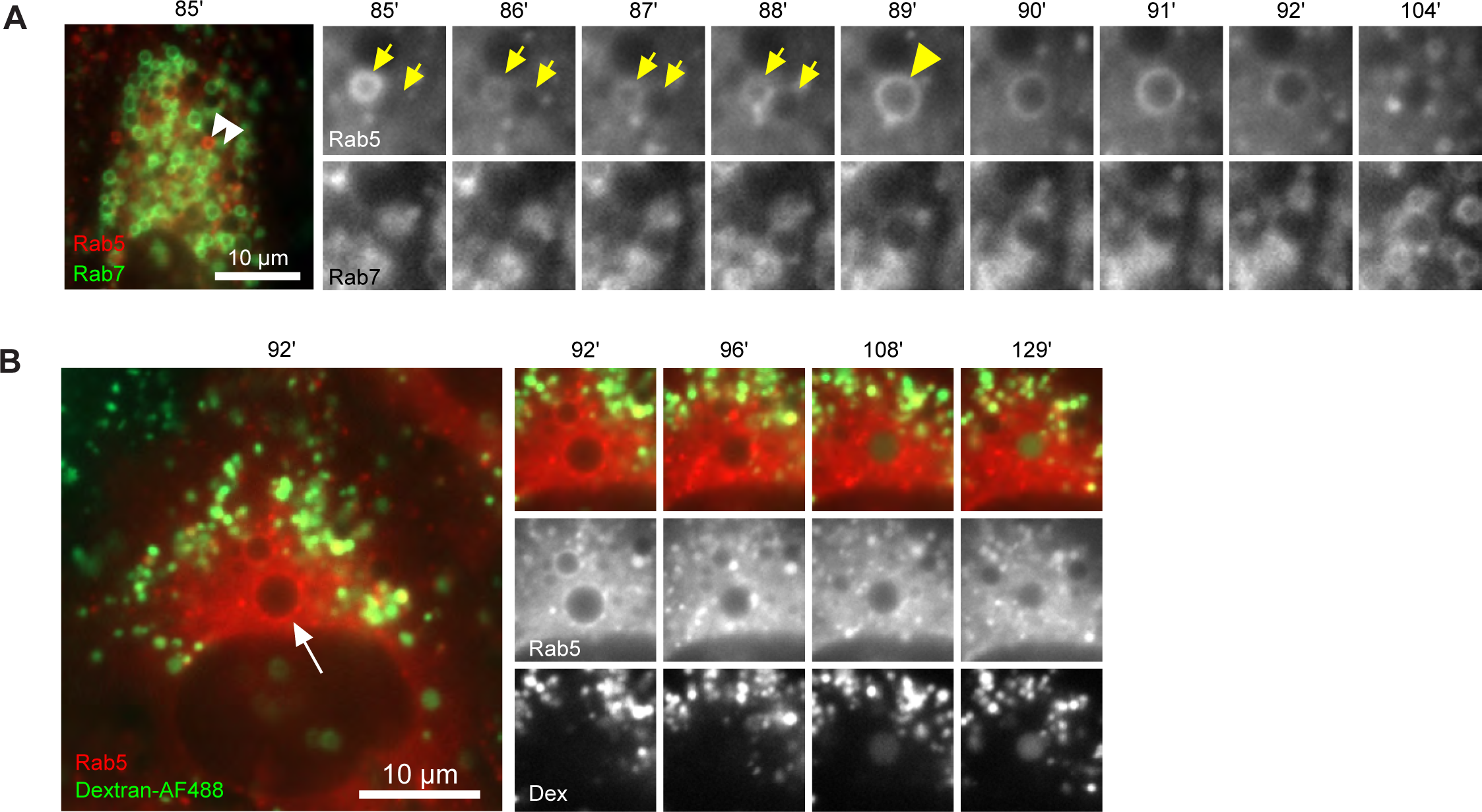
Enlarged compartments interact with early endosomes and acquire intraluminal content from the endolysosomal pathway. Nigericin was added to HeLa cells, stably expressing mApple-Rab5 and GFP-Rab7 (A) or mApple-Rab5 alone (B), for 20 min and washed away, and cells were imaged by tome-lapse microscopy. Recovery times are specified relative to removal of nigericin. (A) Example to show an enlarged compartment devoid of Rab5 fusing with Rab5-positive endosome to subsequently undergo Rab conversion. (B) Example to show an enlarged Rab5-positive endosome later acquiring Dextran-AF488, which had been pulse-chased into the endolysosomal pathway (4h pulse, 1.5 h chase) prior to nigericin treatment. Recovery times are specified relative to removal of nigericin.

**Figure 1, Figure Supplement 2.**
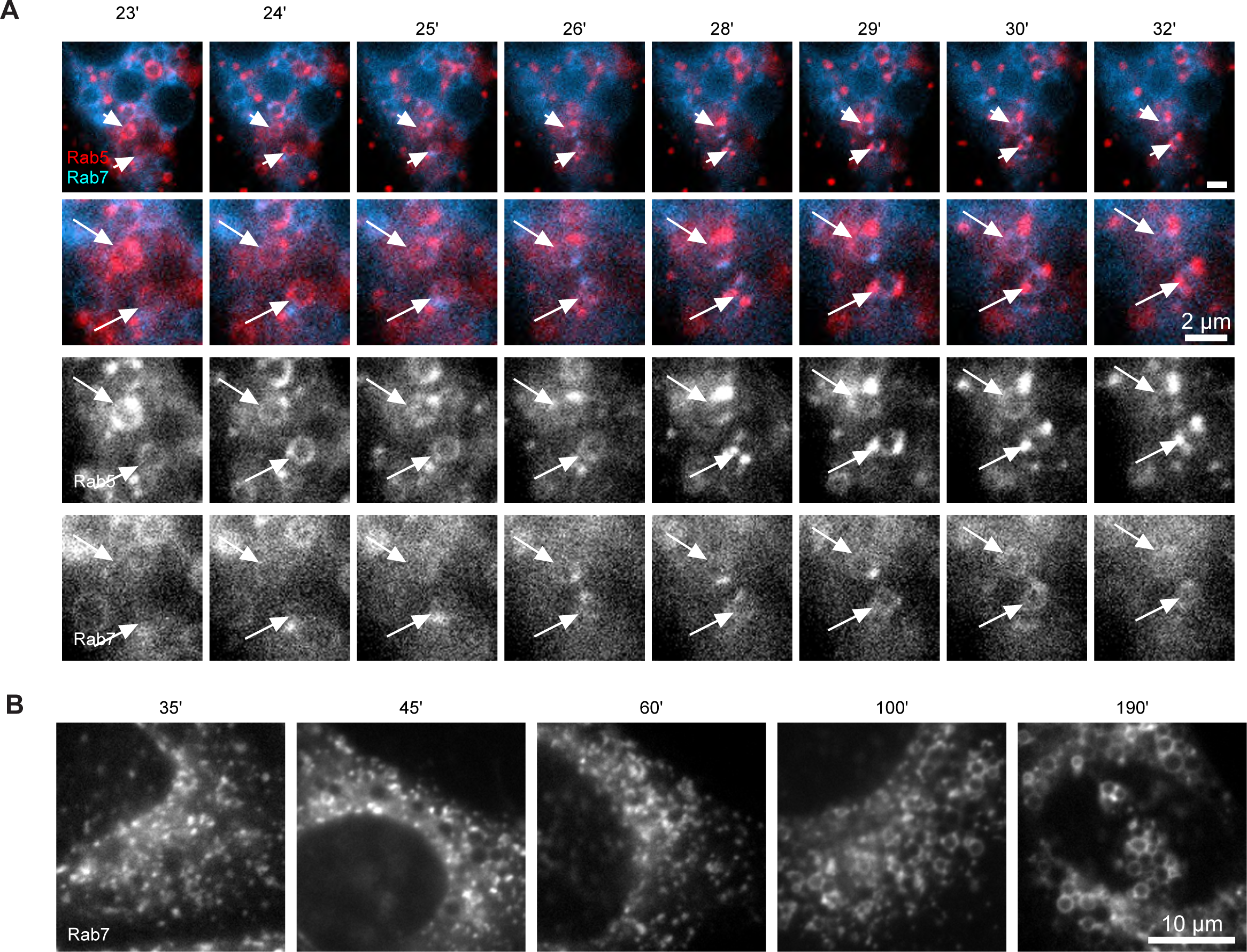
Enlarged endosomes that undergo Rab conversion can be induced by different treatments. (A) HeLa cells stably expressing mApple-Rab5 and transiently transfected with BFP-Rab7 were treated for 4 h with ammonium chloride, washed and allowed to recover for 10 min prior to imaging. Arrows point to two Rab5-positive endosomes, which subsequently acquire Rab7. (B) HeLa cells stably expressing mApple-Rab7 were treated for 20 min with monensin, washed and imaged at specified times after the wash. Representative images of cells show accumulation of enlarged Rab7-positive endosomes at later time points.

**Figure 1, Figure Supplement 3.**
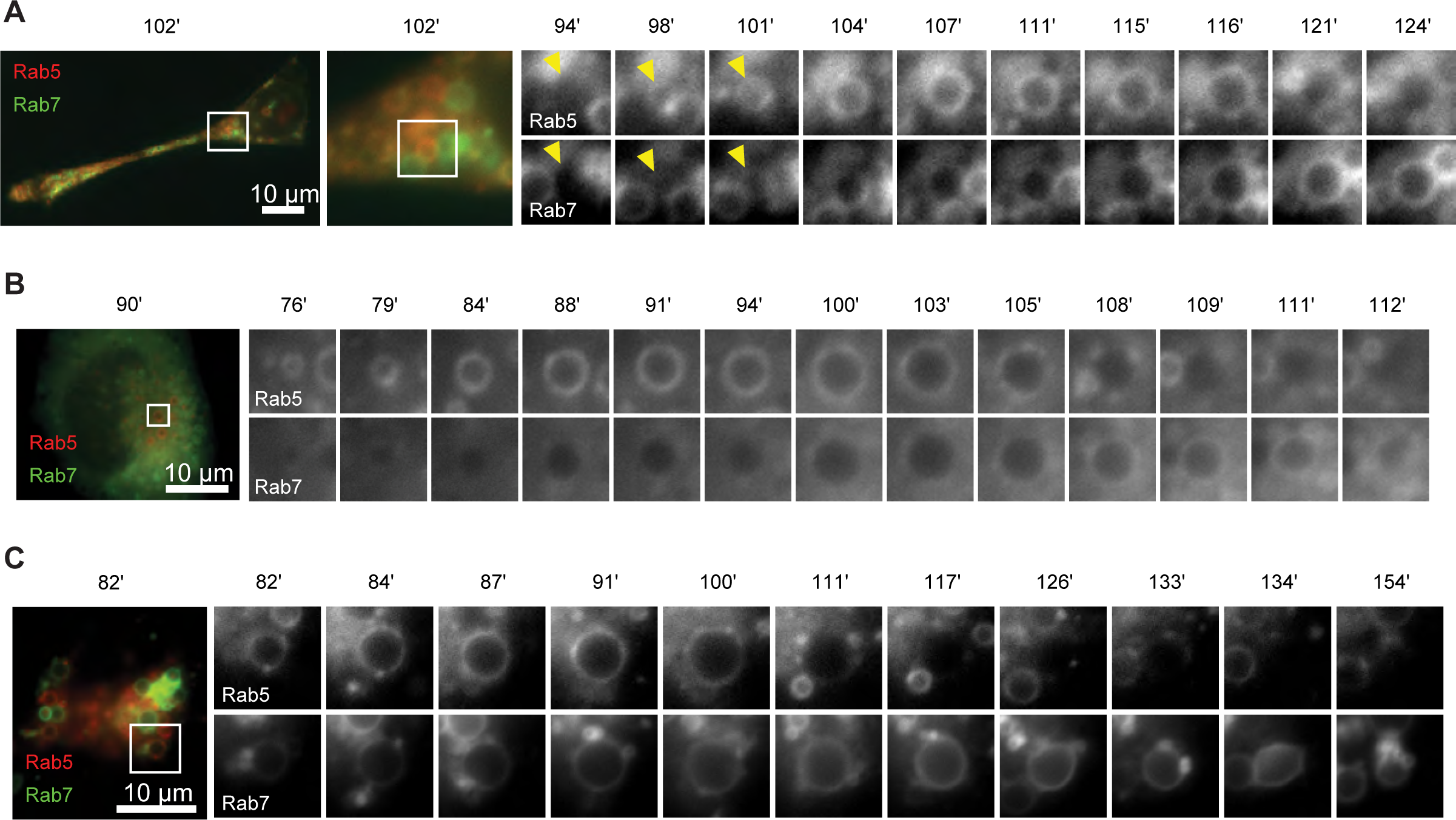
Enlarged endosomes that undergo Rab conversion can be induced in different cell types. Cells were transiently transfected with mApple-Rab5 and GFP-Rab7, and treated 20 min with nigericin followed by recovery and time-lapse microscopy. (A) HEK293 cells, (B) Neuro2A cells, (C) Cos-1 cells. Images were selected to show Rab conversion in the enlarged endosomes.

**Figure 2, Figure Supplement 1.**
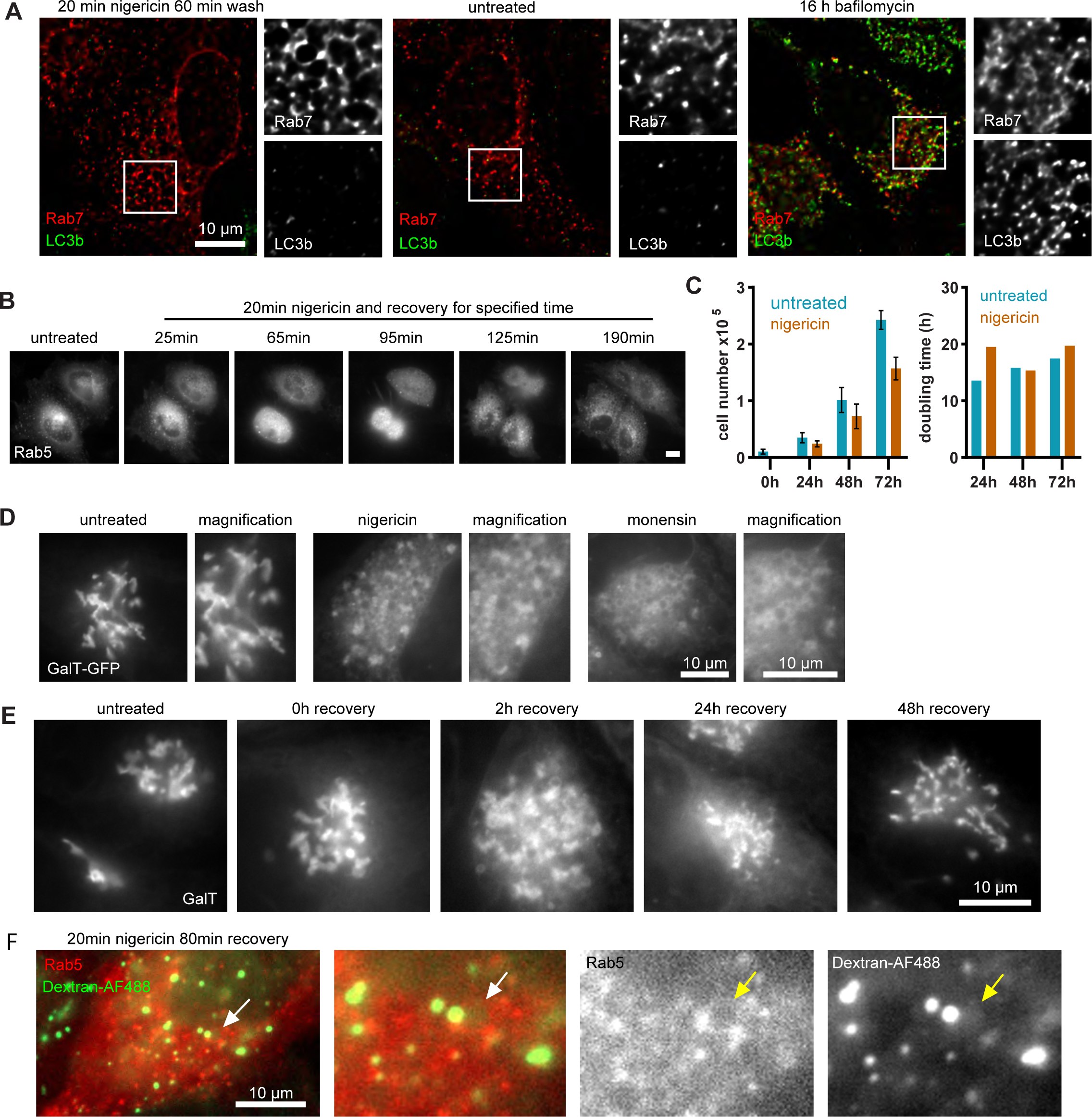
Cell growth is unaffected by nigericin treatment, which induces reversible Golgi vesiculation. HeLa cells stably expressing mApple-Rab5 (A,B,F) or GalT-GFP (C-E) were treated for 20 min with 10 µM nigericin or 5 µM monensin, or left untreated. (A) Images of cells stained with LC3b antibody to show absence of autophagy induction or presence of LC3B at the enlarged endosomes of nigericin-treated cells. Bafilomycin-treated cells are shown here as positive control. (B) Images to show cells dividing shortly after nigericin treatment, with recovery times specified. Scale bar = 10 µm. (C) Cells were plated in 12-well plates at 104 cells per well 24 h prior to nigericin treatment, trypsinised and counted at specified recovery times. Triplicate wells were counted per time point per condition. Actual cells numbers and doubling times are presented. Representative graph of three independent experiments. Source data is available in Figure S4C - Source Data 1. (D Images to reveal extensive formation of GalT-positive enlarged compartments upon either treatment following 2 h recovery. (E) Images to show Golgi vesiculation and return to ribbon morphology within 48 h of recovery from nigericin treatment. (F) Images of cells incubated with 0.5mg/mL Dextran-AF488 for 65min after nigericin washout to reveal endocytosed dextran presence in the enlarged Rab5-positive endosomes.

**Figure 4, Figure Supplement 1.**
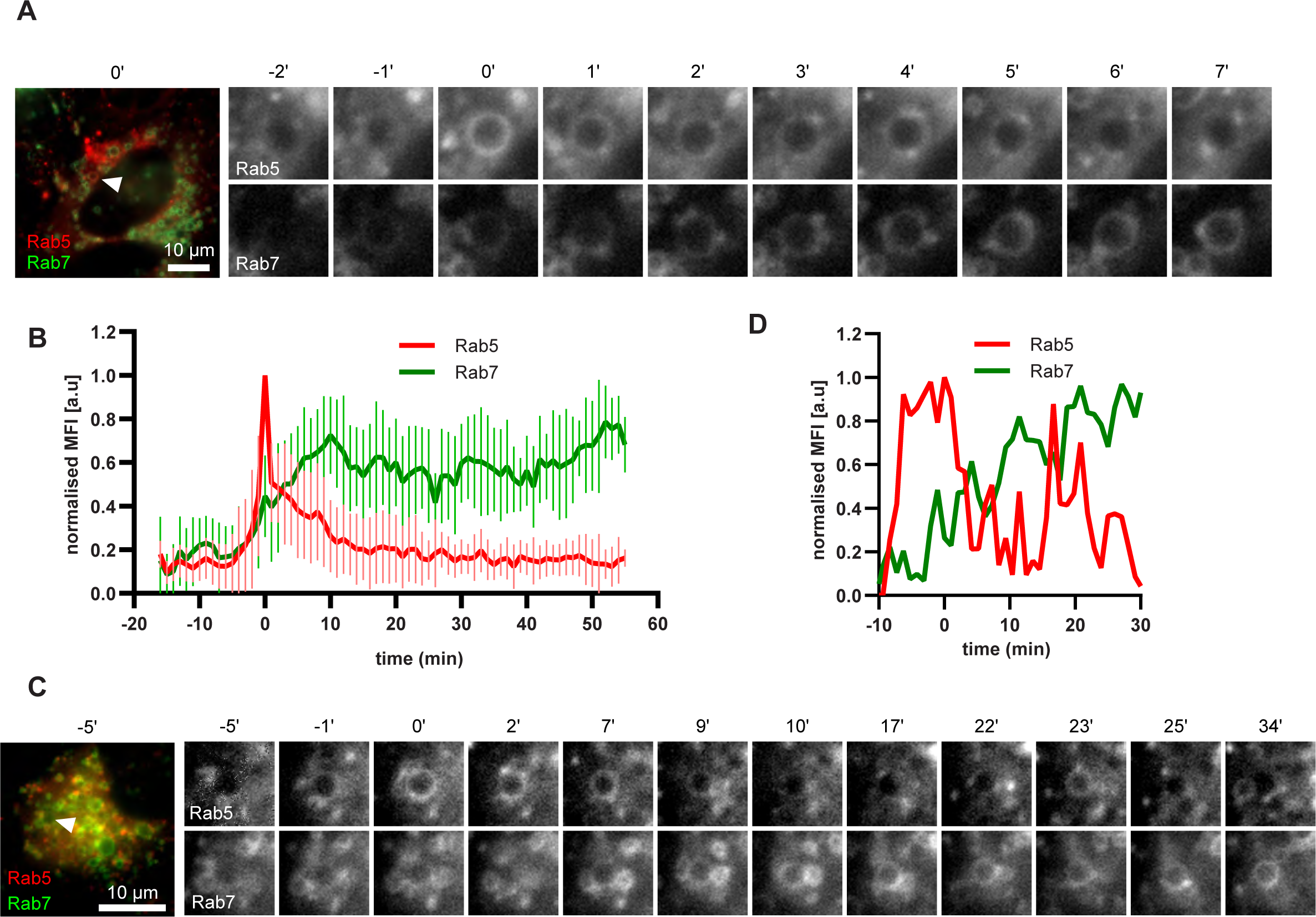
Enlarged endosomes occasionally display multiple Rab5 peaks and form Rab5 subdomains. HeLa cells, stably expressing mApple-Rab5 and GFP-Rab7 were treated for 20 min with nigericin, washed and imaged over 3 h, as described in Figure 4A. (A) Time-lapse images of an endosome displaying formation of Rab5 subdomains. (B) Averaged Rab5 and Rab7 kinetics of 21 endosomes. Error bars represent standard deviation. Representative graph of three independent experiments. (C-D) Time-lapse images (C) and a corresponding fluorescence quantification plot (D) of an endosome with multiple waves of Rab5 recruitment and a continuous increase in Rab7 levels.

**Figure 5, Figure Supplement 1.**
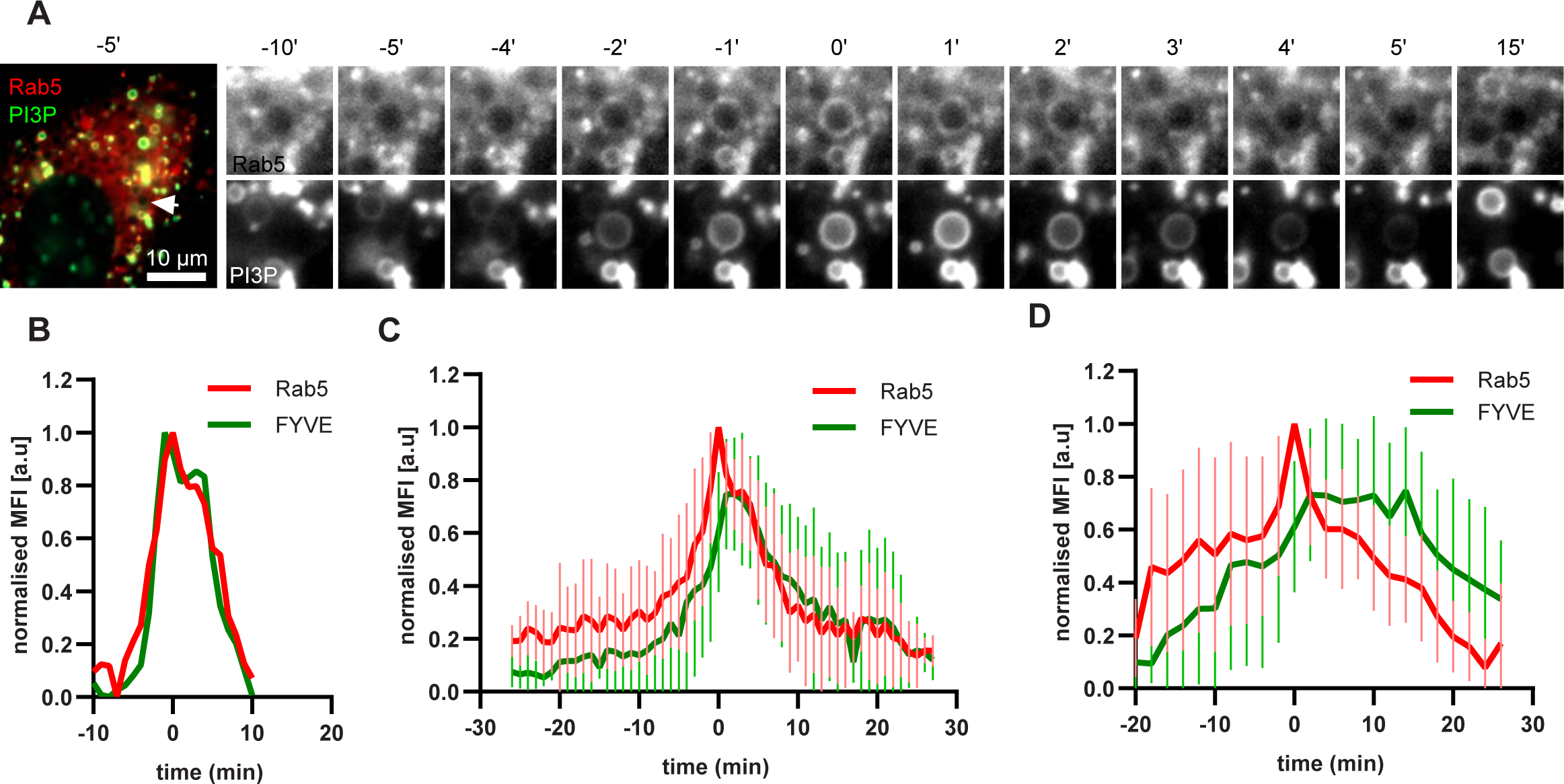
PI(3)P is recruited to endosomes concomitantly with Rab5. HeLa cells, stably expressing mApple-Rab5 and transiently transfected with the PI(3)P marker, GFP-FYVE, were treated for 20 min with nigericin, washed and imaged over 3 h, as described in Figure 4A. (A) Time-lapse images of a representative endosome to show transient Rab5 recruitment accompanied by PI(3)P. (B) Corresponding graph of normalised mean fluorescence intensity of Rab5 and FYVE at the rim of the endosome in (A) over the time the endosome was detectable. (C) Averaged Rab5 and PI(3)P kinetics of 16 endosomes. Error bars represent standard deviation. (D) Averaged Rab5 and PI(3)P kinetics of 15 endosomes. Error bars represent standard deviation. (C) and (D) represent two independent experiments of a total of three.

**Figure 6, Figure Supplement 1.**
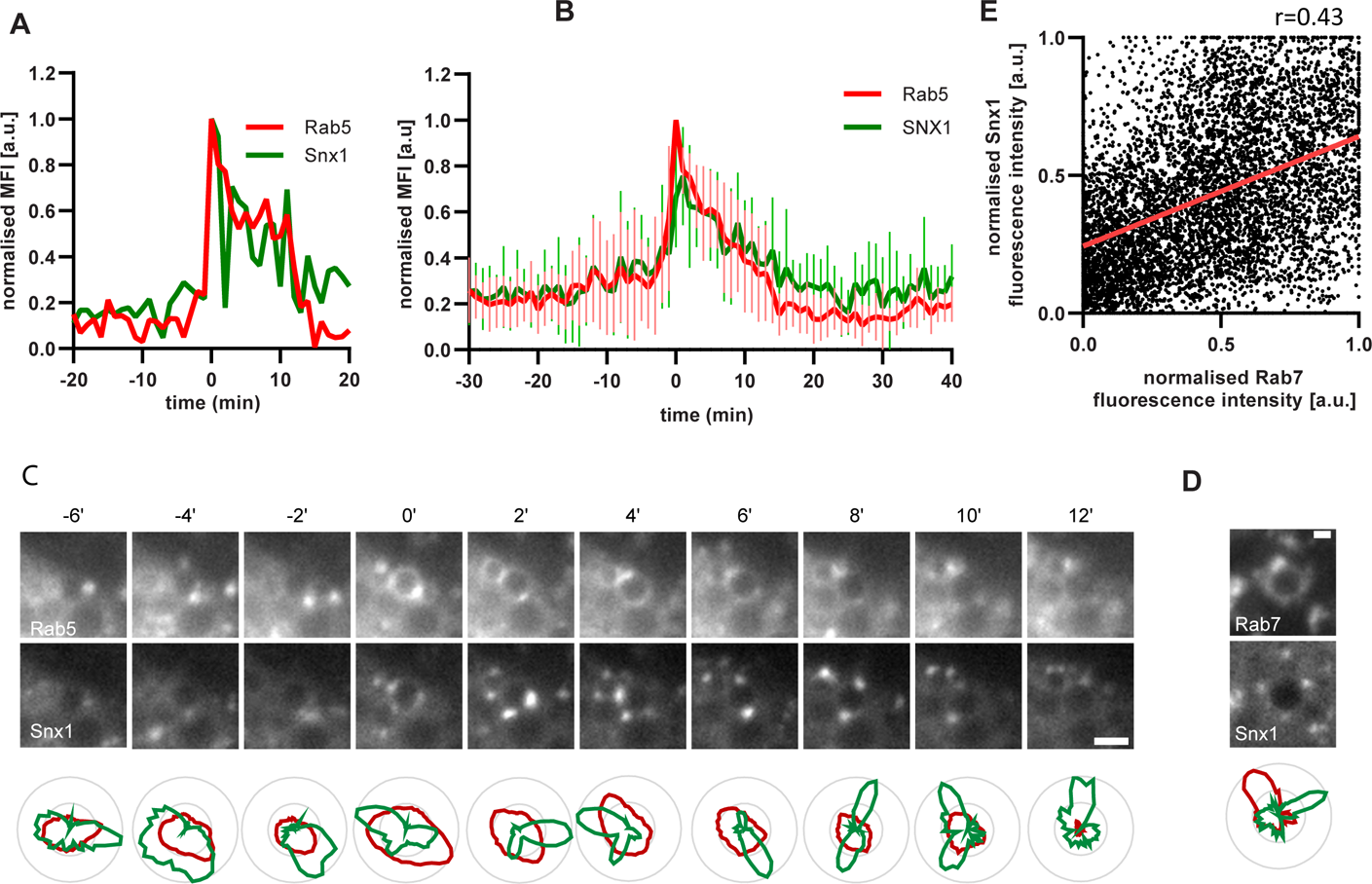
Dynamic Snx1 recruitment suggest active sorting at the enlarged endosome. Nigericin was added to HeLa cells for 20 min and washed away, and cells were imaged by tisme-lapse microscopy, as described in Figure 4A. (A-E) Cells stably expressing mApple-Rab5 (A-C) or mApple-Rab7 (D-E) and transiently transfected with the Snx1-GFP. (A) Example graph of normalised mean fluorescence intensity (MFI) of Rab5 and Snx1 at the rim of the endosome over the time the endosome was detectable. (B) Averaged Rab5 and Snx1 kinetics of 14 endosomes. Error bars represent standard deviation. Representative graph of three independent experiments. Numerical data for all quantified endosomes is available in Figure 6C - Source Data 1. (C) Images and corresponding line profiles of normalised fluorescence intensity of Rab5 and Snx1 along the rim of the maturing endosome at consecutive time points. Rab5 was adjusted for a single maximum and minimum values during the recorded kinetic to highlight its overall signal increase. Snx1 was adjusted for max and min values for each time point to highlight the dynamic nature of Snx1 subdomains. Scale bar = 2 µm. Numerical data for all quantified endosomes is available in Figure S7C - Source Data 1. (D) Images of Rab7 and Snx1 at an enlarged endosome and a corresponding line profile of normalised fluorescence intensity along the rim to show co-existence as well as independence of subdomains of the two markers. Scale bar = 1 µm. (E) Correlation plot of normalised fluorescence intensity of Rab7 and Snx1 as measured in (E) for 3 endosomes for a total of 96 time points, and a corresponding regression line. Pearson’s correlation r=0.43. Numerical data for all quantified endosomes is available in Figure S7E - Source Data 1.

**Figure 8, Figure Supplement 1.**
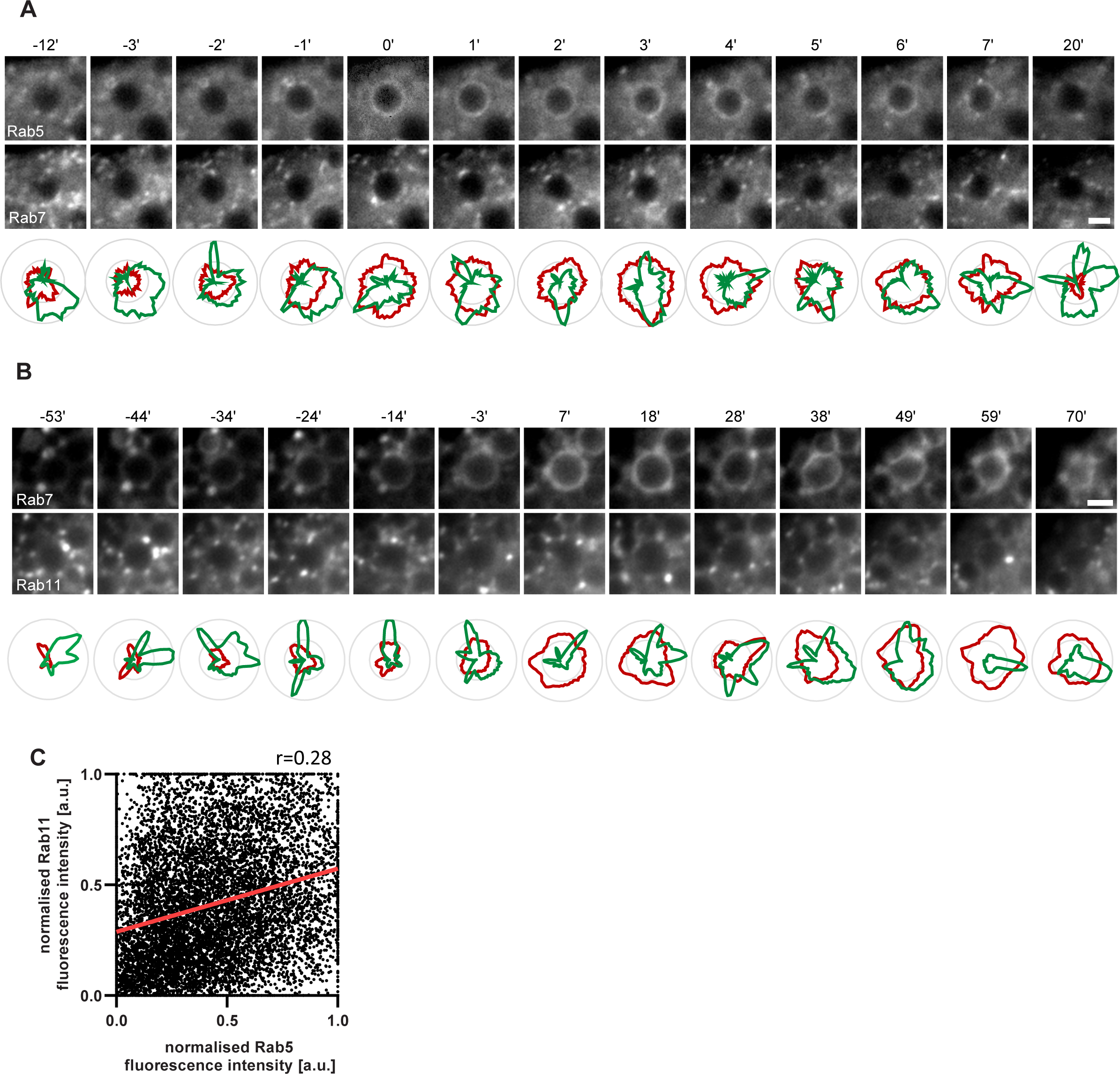
Rab11 interacts with the maturing endosome independently of Rab5 or Rab7. HeLa cells, stably expressing mApple-Rab5 (A,C) or mApple-Rab7 (B) and transiently transfected with GFP-Rab11, were treated for 20 min with nigericin, washed and imaged over 3 h, as described in Figure 4A. (A,B) Images and corresponding line profiles of normalised fluorescence intensity of Rab5 (A) or Rab7 (B) and Rab11 along the rim of the maturing endosome at consecutive time points. Rab5/Rab7 was adjusted for a single maximum and minimum values during the recorded kinetic to highlight its overall signal increase. Rab11 was adjusted for maximum and minimum values for each time point to highlight the dynamic nature of Rab11 interactions with the maturing endosome. Scale bar = 2 µm. (C) Correlation plot of normalised fluorescence intensity of Rab5 and Rab11 as measured in Figure 7D for 14 endosomes for a total of 193 time points, and a corresponding regression line. Pearson’s correlation r=0.28. Numerical data for all analysed endosomes is available in Figure 8D and S8 - Source Data 1 and Figure 8H and S8 - Source Data 1.

**Figure 9, Figure Supplement 1.**
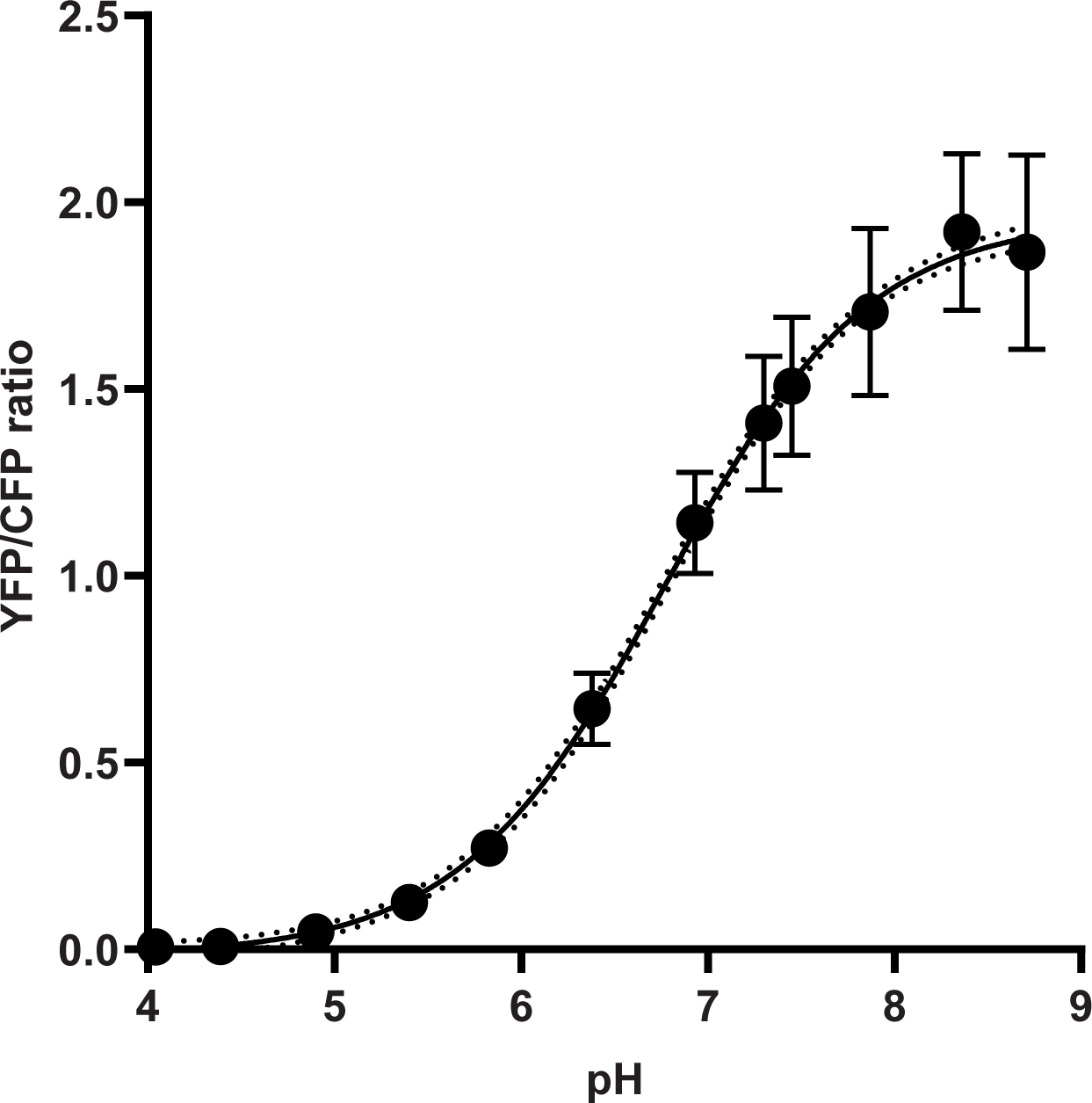
GalT-pHlemon sensor responds to pH changes in a sigmoidal dose-response manner. HeLa cells were transiently transfected with the ratiometric pH sensor, GalT-pHlemon. Graph to show response of GalT-pHlemon sensor to pH 4.0-9.0 range as displayed by YFP/CFP ratio measurements in cells incubated with calibration buffers of specified pH values as well as the interpolation of sigmoidal dose-response curve. Numerical data for all quantified Golgi ROIs is available in Figure S9 - Source Data 1.

**Figure 11, Figure Supplement 2.**
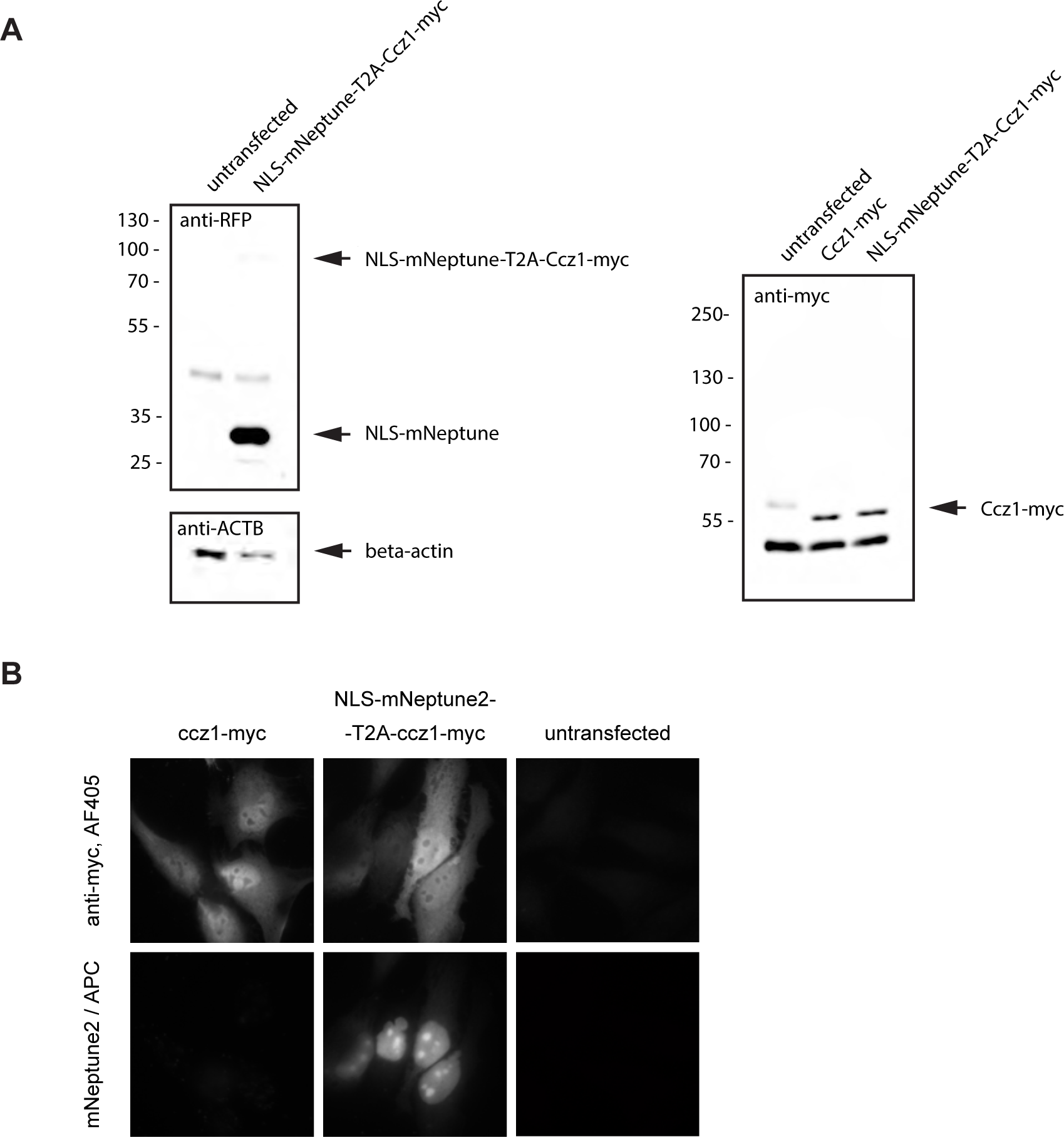
Validation and characterisation of Ccz1 the rescue construct. (A) Western blot of the ccz1 rescue construct expressing NLS-mNeptune2-T2A-Ccz1-myc to show predominant production of two smaller products, NLS-mNeptune2 and ccz1-myc, as visualised by RFP and myc antibodies, respectively. Original uncropped and unformated images are available in Figure S11A - Source Data 1. (B) Immunofluorescence images of NLS-mNeptune2-T2A-Ccz1-myc transiently transfected into HeLa cells to reveal nuclear distribution of the mNeptune2 signal and cytosolic distribution of Ccz1-myc.

**Figure 11, Figure Supplement 1.**
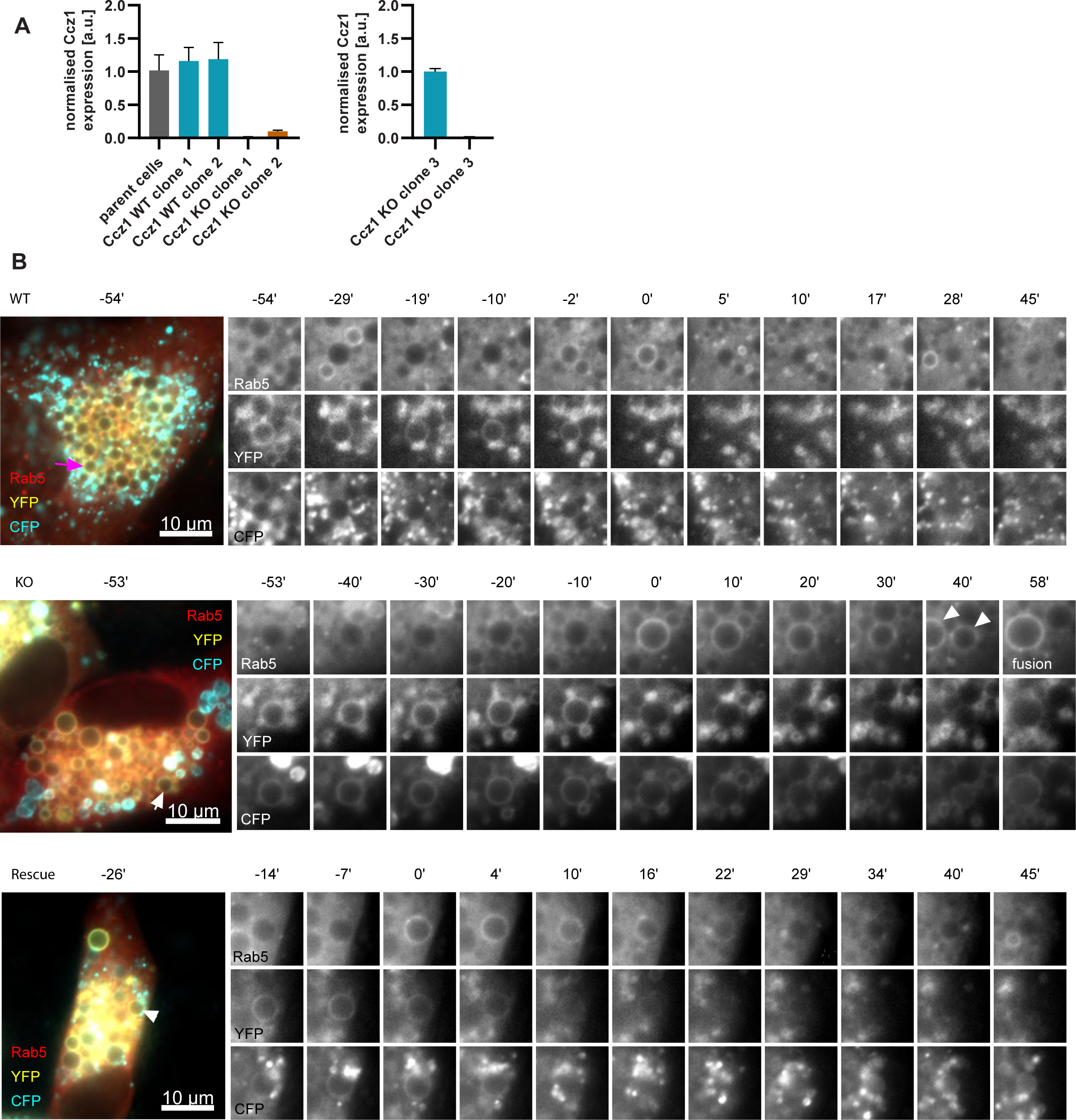
Validation and characterisation of Ccz1 knockout cell lines. (A) Ccz1 mRNA expression levels in HeLa cell lines with wild-type (WT) Ccz1 and knocked-out Ccz1 (KO), three clones each, as measured by qRT-PCR and normalised for acin. Raw RT-PCR data is available in Figure S10A - Source Data 1. (B) HeLa cell lines with wild-type (WT) Ccz1 and knocked-out Ccz1 (KO) were transiently transfected with mAp-ple-Rab5 and GalT-pHlemon. Ccz1 expression plasmid was co-transfected for 72 h for rescue experiments. Time-lapse images of representative endosomes, corresponding to those quantified in Figure 11B, to show acidification in endosomes recruiting Rab5 in the three cell types.

**Figure 12, Figure Supplement 1.**
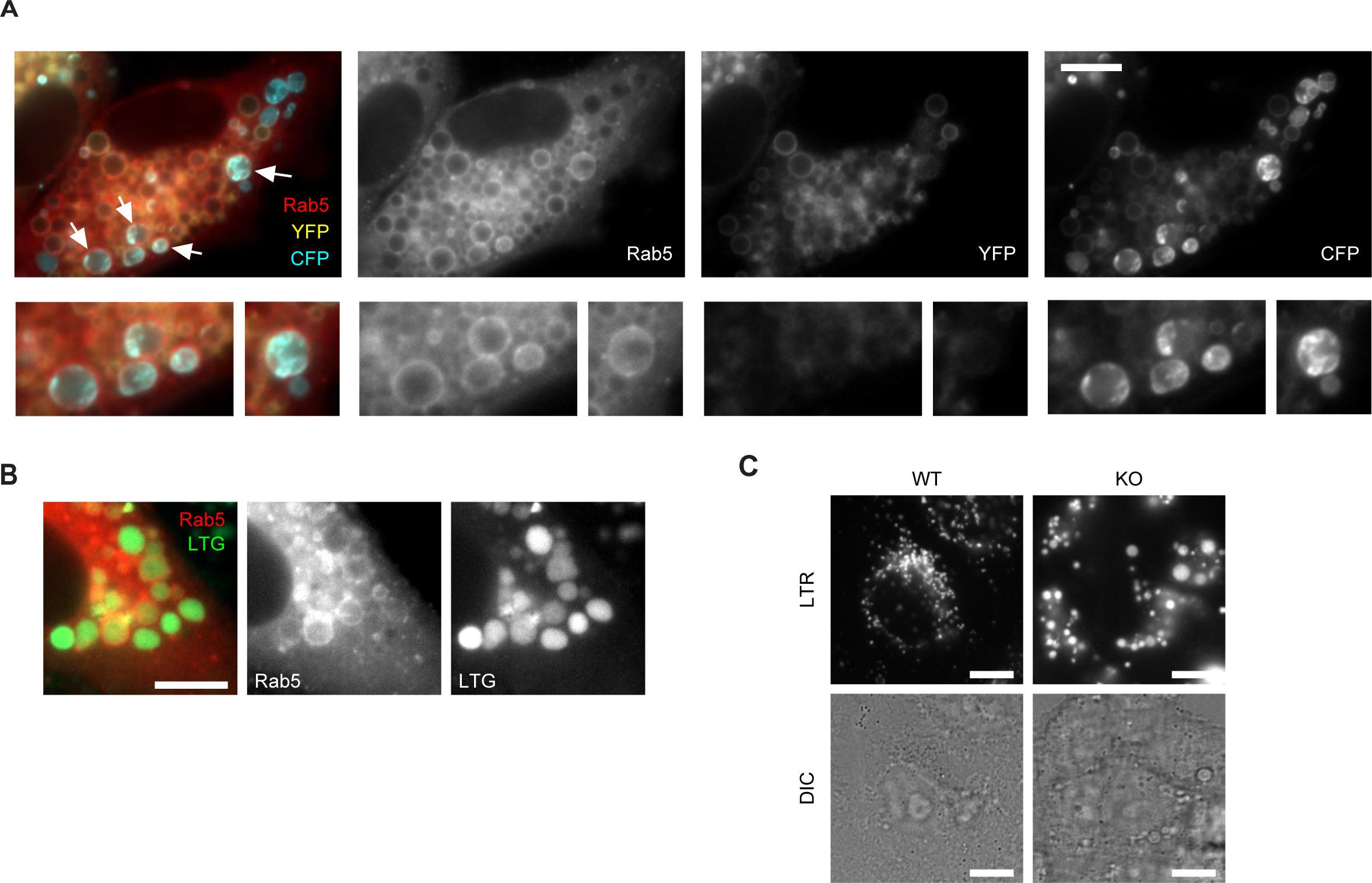
Characterisation of Ccz1 knockout cell lines. (A) Images of Ccz1 KO cells pre-treated with nigericin, recorded at 150 min recovery, showing Rab5-positive acidified hybrid compartments (arrows). Cells were transiently transfected with mApple-Rab5 and GalT-pHlemon. Scale bar = 10 µm. (B) Images of Ccz1 KO cells pre-treated with nigericin, recorded at 24 h recovery, showing Rab5-positive acidified hybrid compartments, as visualised with Lysotracker Green (LTG). Cells were transiently transfected with mAp-ple-Rab5. Scale bar = 10 µm. (C) Images of untreated Ccz1 KO cells stained with Lysotracker Red (LTR) to reveal large, acidified compartments. Scale bar = 10 µm.

## References

1. Balderhaar, H.J., and C. Ungermann. 2013. CORVET and HOPS tethering complexes - coordinators of endosome and lysosome fusion. J Cell Sci. 126:1307–1316.

2. Beuret, N., F. Hasler, C. Prescianotto-Baschong, J. Birk, J. Rutishauser, and M. Spiess. 2017. Amyloid-like aggregation of provasopressin in diabetes insipidus and secretory granule sorting. BMC Biol. 15:5.

3. Bucci, C., P. Thomsen, P. Nicoziani, J. McCarthy, and B. van Deurs. 2000. Rab7: a key to lysosome biogenesis. Mol Biol Cell. 11:467–480.

4. Burgstaller, S., H. Bischof, T. Gensch, S. Stryeck, B. Gottschalk, J. Ramadani-Muja, E. Eroglu, R. Rost, S. Balfanz, A. Baumann, M. Waldeck-Weiermair, J.C. Hay, T. Madl, W.F. Graier, and R. Malli. 2019. pH-Lemon, a Fluorescent Protein-Based pH Reporter for Acidic Compartments. ACS Sens. 4:883–891.

5. Casey, J.R., S. Grinstein, and J. Orlowski. 2010. Sensors and regulators of intracellular pH. Nat Rev Mol Cell Biol. 11:50–61.

6. Choudhury, A., M. Dominguez, V. Puri, D.K. Sharma, K. Narita, C.L. Wheatley, D.L. Marks, and R.E. Pagano. 2002. Rab proteins mediate Golgi transport of caveola-internalized glycosphingolipids and correct lipid trafficking in Niemann-Pick C cells. J Clin Invest. 109:1541–1550.

7. Compton, L.M., O.C. Ikonomov, D. Sbrissa, P. Garg, and A. Shisheva. 2016. Active vacuolar H+ ATPase and functional cycle of Rab5 are required for the vacuolation defect triggered by PtdIns(3,5)P2 loss under PIKfyve or Vps34 deficiency. Am J Physiol Cell Physiol. 311:C366–377.

8. De Luca, M., and C. Bucci. 2014. A new V-ATPase regulatory mechanism mediated by the Rab interacting lysosomal protein (RILP). Commun Integr Biol. 7.

9. De Luca, M., L. Cogli, C. Progida, V. Nisi, R. Pascolutti, S. Sigismund, P.P. Di Fiore, and C. Bucci. 2014. RILP regulates vacuolar ATPase through interaction with the V1G1 subunit. J Cell Sci. 127:2697–2708.

10. Del Conte-Zerial, P., L. Brusch, J.C. Rink, C. Collinet, Y. Kalaidzidis, M. Zerial, and A. Deutsch. 2008. Membrane identity and GTPase cascades regulated by toggle and cut-out switches. Mol Syst Biol. 4:206.

11. Dove, S.K., K. Dong, T. Kobayashi, F.K. Williams, and R.H. Michell. 2009. Phosphatidylinositol 3,5-bisphosphate and Fab1p/PIKfyve underPPIn endo-lysosome function. Biochem J. 419:1–13.

12. Falcon-Perez, J.M., R. Nazarian, C. Sabatti, and E.C. Dell’Angelica. 2005. Distribution and dynamics of Lamp1-containing endocytic organelles in fibroblasts deficient in BLOC-3. J Cell Sci. 118:5243–5255.

13. Gillooly, D.J., I.C. Morrow, M. Lindsay, R. Gould, N.J. Bryant, J.M. Gaullier, R.G. Parton, and H. Stenmark. 2000. Localization of phosphatidylinositol 3-phosphate in yeast and mammalian cells. EMBO J. 19:4577–4588.

14. Guerra, F., and C. Bucci. 2016. Multiple Roles of the Small GTPase Rab7. Cells. 5.

15. Hsu, F., F. Hu, and Y. Mao. 2015. Spatiotemporal control of phosphatidylinositol 4-phosphate by Sac2 regulates endocytic recycling. J Cell Biol. 209:97–110.

16. Huotari, J., and A. Helenius. 2011. Endosome maturation. EMBO J. 30:3481–3500.

17. Ledger, P.W., N. Uchida, and M.L. Tanzer. 1980. Immunocytochemical localization of procollagen and fibronectin in human fibroblasts: effects of the monovalent ionophore, monensin. J Cell Biol. 87:663–671.

18. Ma, L., Q. Ouyang, G.C. Werthmann, H.M. Thompson, and E.M. Morrow. 2017. Live-cell Microscopy and Fluorescence-based Measurement of Luminal pH in Intracellular Organelles. Front Cell Dev Biol. 5:71.

19. McDermott, H., and K. Kim. 2015. Molecular dynamics at the endocytic portal and regulations of endocytic and recycling traffics. Eur J Cell Biol. 94:235–248.

20. Merion, M., and W.S. Sly. 1983. The role of intermediate vesicles in the adsorptive endocytosis and transport of ligand to lysosomes by human fibroblasts. J Cell Biol. 96:644–650.

21. Morre, D.J., W.F. Boss, H. Grimes, and H.H. Mollenhauer. 1983. Kinetics of Golgi apparatus membrane flux following monensin treatment of embryogenic carrot cells. Eur J Cell Biol. 30:25–32.

22. Nagano, M., J.Y. Toshima, D.E. Siekhaus, and J. Toshima. 2019. Rab5-mediated endosome formation is regulated at the trans-Golgi network. Commun Biol. 2:419.

23. Naufer, A., V.E.B. Hipolito, S. Ganesan, A. Prashar, V. Zaremberg, R.J. Botelho, and M.R. Terebiznik. 2018. pH of endophagosomes controls association of their membranes with Vps34 and PtdIns(3)P levels. J Cell Biol. 217:329–346.

24. Nordmann, M., M. Cabrera, A. Perz, C. Brocker, C. Ostrowicz, S. Engelbrecht-Vandre, and C. Ungermann. 2010. The Mon1-Ccz1 complex is the GEF of the late endosomal Rab7 homolog Ypt7. Curr Biol. 20:1654–1659.

25. Peter, B.J., H.M. Kent, I.G. Mills, Y. Vallis, P.J. Butler, P.R. Evans, and H.T. McMahon. 2004. BAR domains as sensors of membrane curvature: the amphiphysin BAR structure. Science. 303:495–499.

26. Podinovskaia, M., W. Lee, S. Caldwell, and D.G. Russell. 2013. Infection of macrophages with Mycobacterium tuberculosis induces global modifications to phagosomal function. Cell Microbiol. 15:843–859.

27. Podinovskaia, M., and A. Spang. 2018. The Endosomal Network: Mediators and Regulators of Endosome Maturation. Prog Mol Subcell Biol. 57:1–38.

28. Poteryaev, D., S. Datta, K. Ackema, M. Zerial, and A. Spang. 2010. Identification of the switch in early-to-late endosome transition. Cell. 141:497–508.

29. Poteryaev, D., H. Fares, B. Bowerman, and A. Spang. 2007. Caenorhabditis elegans SAND-1 is essential for RAB-7 function in endosomal traffic. EMBO J. 26:301–312.

30. Ran, F.A., P.D. Hsu, J. Wright, V. Agarwala, D.A. Scott, and F. Zhang. 2013. Genome engineering using the CRISPR-Cas9 system. Nat Protoc. 8:2281–2308.

31. Rink, J., E. Ghigo, Y. Kalaidzidis, and M. Zerial. 2005. Rab conversion as a mechanism of progression from early to late endosomes. Cell. 122:735–749.

32. Rojas, R., T. van Vlijmen, G.A. Mardones, Y. Prabhu, A.L. Rojas, S. Mohammed, A.J. Heck, G. Raposo, P. van der Sluijs, and J.S. Bonifacino. 2008. Regulation of retromer recruitment to endosomes by sequential action of Rab5 and Rab7. J Cell Biol. 183:513–526.

33. Schink, K.O., C. Raiborg, and H. Stenmark. 2013. Phosphatidylinositol 3-phosphate, a lipid that regulates membrane dynamics, protein sorting and cell signalling. Bioessays. 35:900–912.

34. Skjeldal, F.M., L.H. Haugen, D. Mateus, D.M. Frei, A.V. Rodseth, X. Hu, and O. Bakke. 2021. De novo formation of early endosomes during Rab5-to-Rab7a transition. J Cell Sci. 134.

35. Solinger, J.A., H.O. Rashid, C. Prescianotto-Baschong, and A. Spang. 2020. FERARI is required for Rab11-dependent endocytic recycling. Nat Cell Biol. 22:213–224.

36. Solinger, J.A., and A. Spang. 2013. Tethering complexes in the endocytic pathway: CORVET and HOPS. FEBS J. 280:2743–2757.

37. Spang, A. 2016. Membrane Tethering Complexes in the Endosomal System. Front Cell Dev Biol. 4:35.

38. van den Boomen, D.J.H., A. Sienkiewicz, I. Berlin, M.L.M. Jongsma, D.M. van Elsland, J.P. Luzio, J.J.C. Neefjes, and P.J. Lehner. 2020. A trimeric Rab7 GEF controls NPC1-dependent lysosomal cholesterol export. Nat Commun. 11:5559.

39. van der Schaar, H.M., M.J. Rust, C. Chen, H. van der Ende-Metselaar, J. Wilschut, X. Zhuang, and J.M. Smit. 2008. Dissecting the cell entry pathway of dengue virus by single-particle tracking in living cells. PLoS Pathog. 4:e1000244.

40. van Weering, J.R., P. Verkade, and P.J. Cullen. 2012. SNX-BAR-mediated endosome tubulation is co-ordinated with endosome maturation. Traffic. 13:94–107.

41. Wu, Y., C. Boulogne, S. Carle, M. Podinovskaia, H. Barth, A. Spang, J.C. Cintrat, D. Gillet, and J. Barbier. 2020. Regulation of endo-lysosomal pathway and autophagic flux by broad-spectrum antipathogen inhibitor ABMA. FEBS J. 287:3184–3199.

42. Zerial, M., and H. McBride. 2001. Rab proteins as membrane organizers. Nat Rev Mol Cell Biol. 2:107–117.

